# Blood-brain barrier-traversing multivalent TGFβ trap for anti-cancer immunomodulation of gliomas

**DOI:** 10.64898/2026.10.05.756837

**Authors:** Daiheon Lee, Seung Woo Chung, Byoungjae Kong, Yunxuan Xie, Bokyoung Kang, Jun Yang, Buwei Huang, Divya Rao, Jong Seob Choi, Eun A. Ra, Pavlos Anastasiadis, Gabsang Lee, Deok-Ho Kim, Michael Lim, Jung Soo Suk

## Abstract

Clinical studies have revealed that glioblastoma (GBM) is refractory to immunotherapy, such as immune checkpoint inhibitors (ICIs), due to the immunosuppressive tumor microenvironment (TME) primarily intensified by intratumoral upregulation of transforming growth factor-β (TGFβ). Primary GBM tumor bulks are surgically removed in clinic, but invasive tumor cells infiltrated into surrounding normal brain tissue cannot be eradicated by systemic therapy due to the integral blood-brain barrier (BBB), leading to near-inevitable tumor recurrence. To address these translational challenges, we have engineered a TGFβ-inhibiting protein nanocage comprising human heavy-chain ferritin protein and TGFβ receptor II ectodomain that enables BBB penetration and multivalent TGFβ entrapment, respectively. We demonstrate that the nanocage platform, denoted TβR2-HFt, following systemic administration, traverses the BBB, and extensively modulates immunosuppressive TME, both stromal and immune components, thereby potentiating the therapeutic efficacy of a clinically used ICI in multiple mouse glioma models. Notably, a majority of the glioma-bearing animals that received surgical tumor resection, followed by treatment with TβR2-HFt and the ICI, survived the tumor and resisted tumor rechallenge, providing a potential means to control GBM in a durable manner.

**Teaser:** BBB-traversing multivalent TGFβ trap reverses the immunosuppressive TME to rescue therapeutic efficacy of immune checkpoint blockade against GBM.

## Introduction

Glioblastoma (GBM) cannot be completely eradicated by the current aggressive standard-of-care treatment regimen while highly lethal tumor recurrence is near-inevitable (*1*). Immunotherapy, such as immune checkpoint inhibitors (ICIs), has brought about transformative control over multiple malignancies (*2*), but clinical studies have revealed poor efficacy against GBM (*3, 4*). The reality is primarily attributed to the notoriously immunosuppressive tumor microenvironment (TME) in GBM (*5–7*). Transforming growth factor-β (TGFβ) is a central immunosuppressive driver (*8, 9*), which, via its canonical intracellular SMAD signaling pathway, promotes extracellular matrix (ECM) remodeling and fibrotic programs to restrict immune cell access to tumors (*10–12*). TGFβ also directly suppresses anti-tumor T-cell responses to drive immune evasion and resistance to immunotherapy (*9, 13*). TGFβ inhibition thus has emerged as a viable strategy to reverse the immunosuppressive TME, thereby restoring anti-tumor immunity and rescuing the therapeutic efficacy of immunotherapy (*10, 11*). Multiple preclinical and clinical studies have demonstrated proof-of-concept efficacy of dual TGFβ-immune checkpoint blockade against non-brain tumors (*10, 11, 14–16*), but its efficacy against GBM remained limited in a clinical study (*17*).

A unique challenge to implementing the strategy against GBM is the blood-brain barrier (BBB) that restricts the delivery of systemically administered therapeutics into the brain parenchyma. Of note, only ∼0.1% of systemically administered antibodies can traverse the intact BBB at best (*18*). Although BBB is often disrupted in high-grade gliomas, its barrier function is retained throughout the tumor regions, particularly the invasive tumor margins where tumor cells infiltrate into surrounding normal brain tissue perfused by blood vessels with integral BBB (*19*). Reflecting this reality, systemically administered anti-TGFβ antibody has shown negligible therapeutic performance against an orthotopic mouse glioma, in contrast to its remarkable efficacy in an otherwise identical subcutaneous glioma model (*20*). These prior observations collectively led to our hypothesis that extensive intratumoral TGFβ inhibition and subsequent restoration of ICI efficacy would be achieved by addressing this globally appreciated delivery challenge.

To achieve this goal, we have leveraged human heavy-chain ferritin nanocage (HFt) as a delivery platform primarily due to its inherent ability to traverse the BBB via the transferrin receptor 1 (TfR1)-mediated transcytosis (*21, 22*). Specifically, we have engineered a hybrid nanocage, denoted TβR2-HFt, based on a fusion protein of human heavy-chain ferritin protein (FHC) and TGFβ receptor II ectodomain (TGFβR2ecd), as a TGFβ trapping domain. The nanocages are formed by self-assembly of 24 protein subunits (*23*) to display up to 24 TGFβR2ecd molecules on the nanocage surface, which enables multivalent TGFβ trapping and inhibition. Further, TβR2-HFt is capable of binding to TfR1 upregulated on GBM cell surface (*24*) to prolong its intratumoral retention and immunomodulatory effects. To this end, the overarching goal of this study is to demonstrate that systemically administered TβR2-HFt would traverse the BBB to deter the immunosuppressive TGFβ/SMAD pathway in TME in a multivalent and prolonged manner to restore the therapeutic efficacy of a clinically used ICI against GBM (**Fig. 1**).

**Fig. 1.**
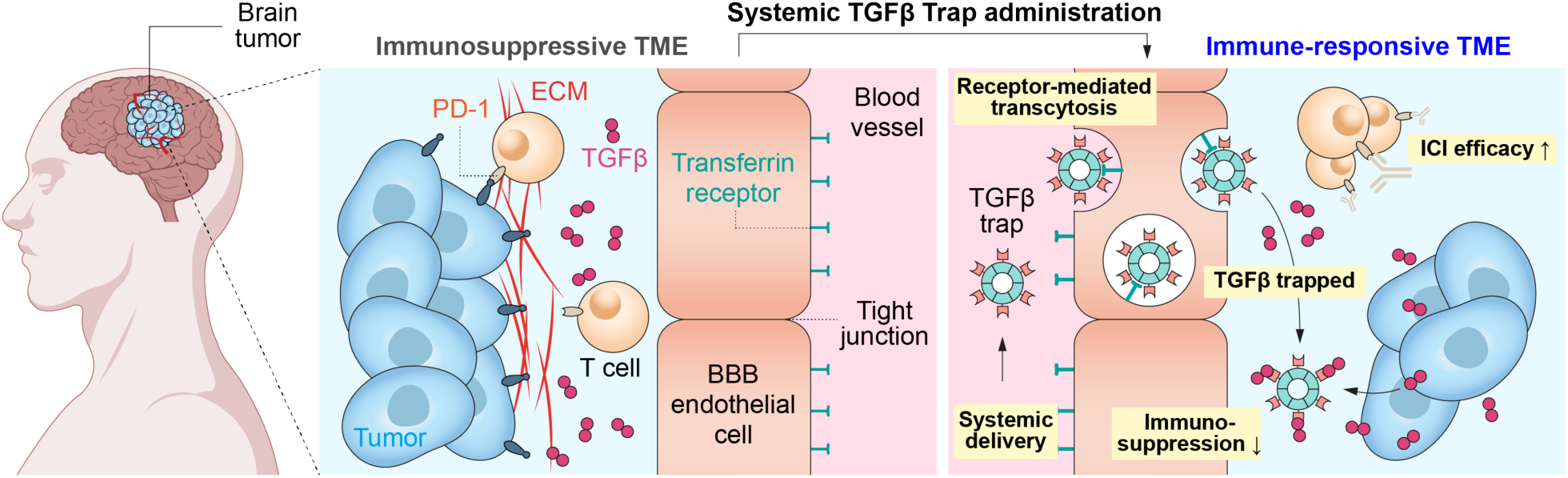
The proposed delivery and therapeutic mechanism of a novel BBB-traversing multivalent TGFβ trap, TβR2-HFt. TβR2-HFt is engineered to traverse the BBB via receptor-mediated transcytosis following systemic administration and subsequently trap TGFβ in a multivalent manner to reverse the immunosuppressive TME to ultimately rescue the therapeutic efficacy of otherwise inefficacious immunotherapy, such as ICIs, against GBM.

## Results

### TGFβ isoforms and their receptors are upregulated and positively correlated with stromal fibrosis and immune suppression in human GBM

To establish target relevance, we first examined the mRNA transcript levels of TGFβ isoforms and receptors in human glioma specimens, in comparison to normal human brain tissues, using the datasets from the Genotyping-Tissue Expression project (GTEx) and The Cancer Genome Atlas (TCGA). We found that the levels of *TGFB1*, *TGFB2*, *TGFB3*, *TGFBR1*, and *TGFBR2* mRNA transcripts were greater in low-grade glioma (LGG) and GBM, more so in GBM, than in normal brains (**Fig. 2a**; normal, *n* = 1,142; LGG, *n* = 509; GBM, *n* = 153). We next analyzed the correlations between the transcript levels of TGFβ isoforms and those of stromal fibrosis and immune suppression markers in GBM, using the datasets from TCGA and Chinese Glioma Genome Atlas (CGGA). Numerical scores for TGFβ, stromal fibrosis, and immune suppression were calculated from the average transcript levels of TGFβ isoforms and key stromal fibrosis (*FN1*, *COL1A1*, *COL5A3*, and *ACTA2*), and immune suppression (*CD274*, *HAVCR2*, *FOXP3*, and *IL2RA*) markers, respectively. Our analysis revealed positive correlations of TGFβ score with stromal fibrosis (r = 0.4746) and immune suppression (r = 0.4743) scores in the TCGA-GBM cohort (**Fig. 2b**). Likewise, TGFβ score was positively correlated with both stromal fibrosis (r = 0.7829) and immune suppression (r = 0.7089) scores in the CGGA-GBM cohort (**Fig. 2c**). We then analyzed the correlations between the transcript levels of each TGFβ isoform and those of an expanded list of individual stromal fibrosis and immune suppression markers. In both TCGA-GBM and CGGA-GBM cohorts, *TGFB1* transcript level consistently exhibited positive correlations with all analyzed markers (**Figs. 2d** and **S1**). In contrast, the correlations with *TGFB2* and *TGFB3* transcript levels varied with individual markers and cohorts (**Figs. 2d** and **S1**).

**Fig. 2.**
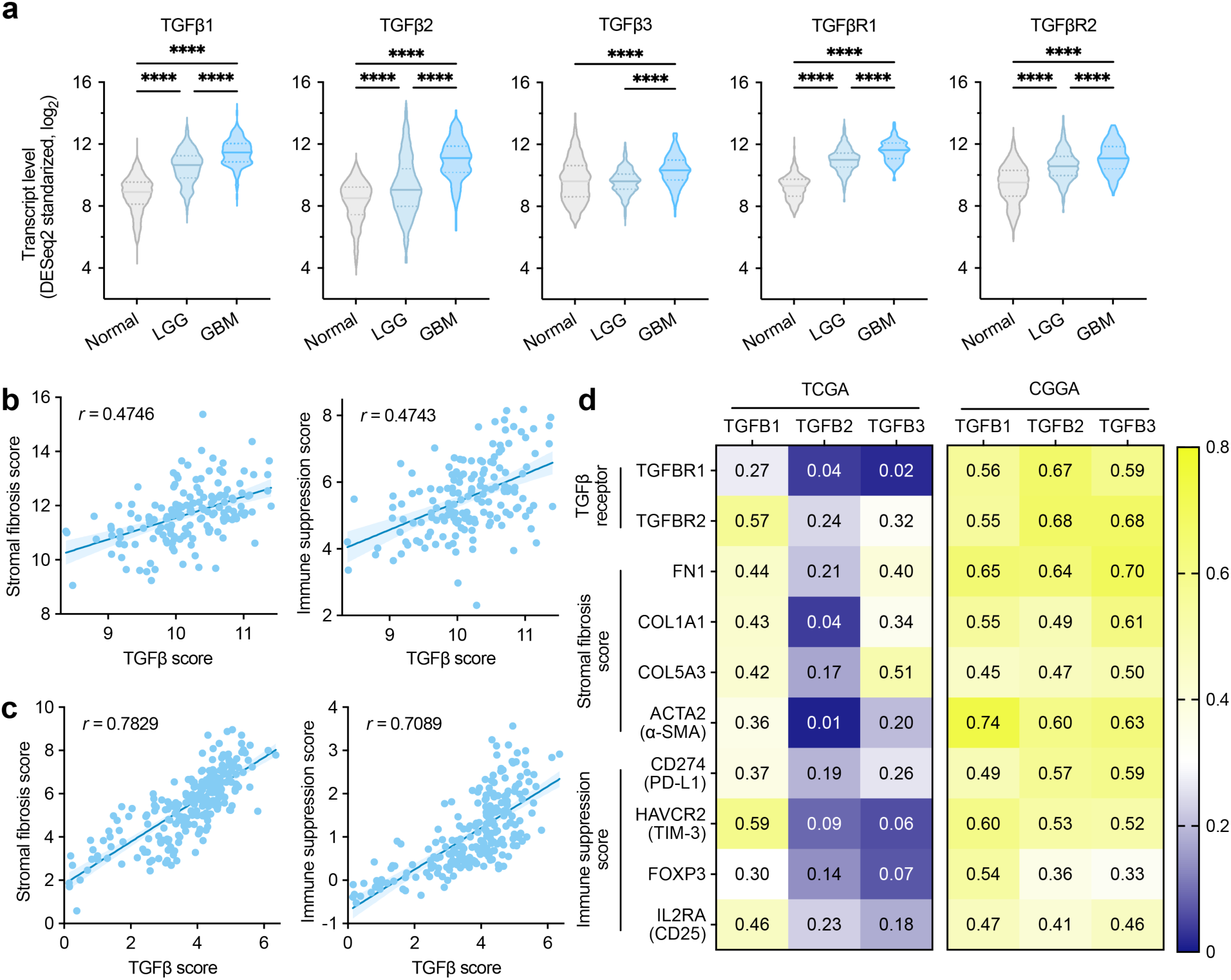
TGFβ is upregulated in GBM and positively correlated with stromal fibrosis and immune suppression in TME. **a**, *TGFB1*, *TGFB2*, *TGFB3*, *TGFBR1*, and *TGFBR2* mRNA transcript levels in human low-grade glioma (LGG) and glioblastoma (GBM) compared to normal brain. The analysis is based on GTEx and TCGA datasets (Normal, n = 1,142; LGG, n = 509; GBM, n = 153). **b**,**c**, Positive correlations between TGFβ and stromal fibrosis or immune suppression scores in TCGA-GBM (**b**) and CGGA-GBM (**c**) cohorts. **d**, Correlations of individual TGFβ isoforms with their receptors or stromal fibrosis and immune suppression markers in TCGA-GBM and CGGA-GBM cohorts. Stromal fibrosis markers: *FN1*, *COL1A1*, *COL5A3*, and *ACTA2*; immune suppression markers: *CD274*, *HAVCR2*, *FOXP3*, *IL2RA*. Statistical significance was determined by one-way ANOVA with Tukey’s post hoc test for **a**. Correlations in **b-d** were assessed by Pearson’s correlation analysis. \*\*\*\**p* < 0.0001.

### TβR2-HFt engineered with a fusion protein of TGFβR2ecd C-terminally fused to FHC exhibits robust TGFβ inhibitory activity

We designed recombinant fusion proteins of FHC and TGFβR2ecd to engineer a hybrid nanocage capable of traversing the BBB and trapping TGFβ in a multivalent manner (**Fig. 3a**). We generated four fusion protein variants with distinct domain orders of FHC and TGFβR2ecd connected via either a flexible or a rigid linker (**Fig. 3b** and **Table S1**). Production of the protein variants was confirmed by SDS-PAGE under reducing condition where the protein bands for the variants with flexible (variant 1 and 3) and rigid (variant 2 and 4) linkers appeared near the predicted molecular weights of 40 and 43 kDa, respectively (**Fig. 3c**). We next screened the ability of four fusion protein nanocage variants to inhibit canonical TGFβ/SMAD signaling pathway, using a reporter cell system that expresses luciferase upon stimulation with recombinant human TGFβ1. The nanocages based on the fusion proteins of TGFβR2ecd fused to the C-terminus of FHC (variants 1 and 2) exhibited robust TGFβ inhibitory activity, but those with the opposite domain order (variants 3 and 4) were unable to inhibit the signaling pathway (**Fig. 3d**). The IC_50_ value of the nanocage with a flexible linker (variant 1; 14.08 pM; 95% CI, 12.30-16.15 pM) was slightly lower (*i.e*., greater TGFβ inhibitory activity) than that of the nanocage with a rigid linker (variant 2; 21.52 pM; 95% CI, 18.13-25.56 pM) (**Fig. 3e**). Thus, the former was selected and designated as TβR2-HFt to be further evaluated. We then compared the physicochemical properties of TβR2-HFt and its parent HFt using size exclusion chromatography (SEC) and dynamic light scattering (DLS). The SEC analysis showed an earlier elution of TβR2-HFt compared to HFt, reflecting the increase in size by the fusion with TGFβR2ecd (**Fig. 3f**). In agreement with the result of SEC analysis, the hydrodynamic diameters of HFt and TβR2-HFt measured by DLS were 14.3 ± 0.1 nm and 29.3 ± 0.3 nm, respectively (**Fig. 3g**).

**Fig. 3.**
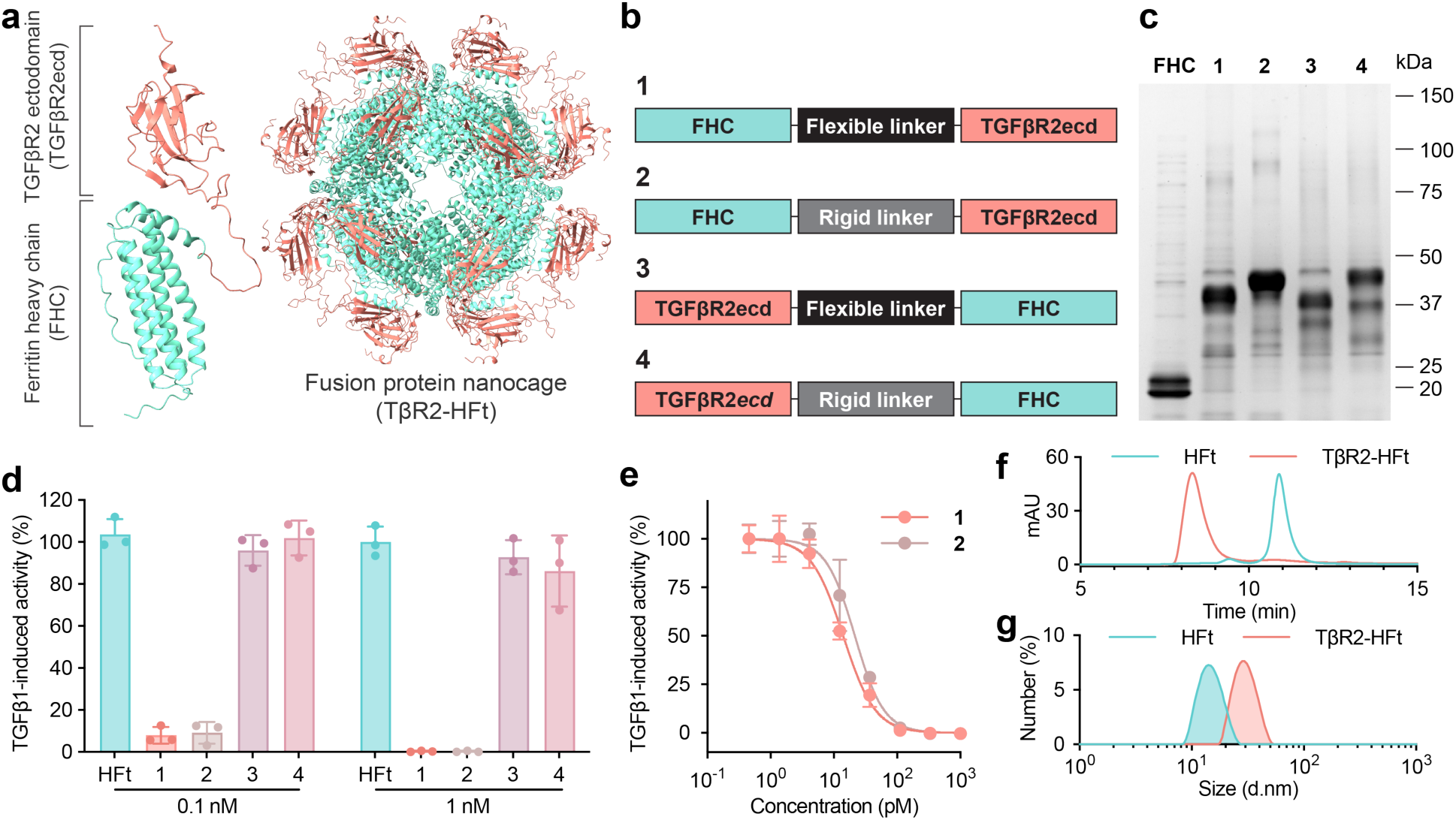
TβR2-HFt engineered with a fusion protein of TGFβR2ecd C-terminally fused to FHC exhibits robust TGFβ inhibitory activity. **a**, Schematics of 3D-modeled structures of a monomeric TGFβR2ecd-FHC subunit (left) and a self-assembled multimeric protein nanocage, TβR2-HFt (right). **b**, Schematic diagrams of cDNA open reading frames of four fusion protein variants with distinct domain orders and linker types. **c**, SDS-PAGE analysis of FHC and fusion protein variants under a reducing condition. **d**, TGFβ inhibitory activity of nanocages based on the fusion protein variants in **b** against recombinant human TGFβ1 in TGFβ/SMAD signaling reporter cells. **e**, Dose-dependent inhibition of TGFβ/SMAD signaling by fusion protein nanocage variants 1 and 2 in the reporter cells. **f**,**g**, Size-exclusion chromatography (**f**) and dynamic light scattering (**g**) analyses of parent HFt and fusion protein nanocage variant 1, TβR2-HFt. Data are presented as mean ± s.d. in **d** and **e**.

### TβR2-HFt shows high-affinity pan-TGFβ binding and markedly enhanced TGFβ inhibitory activity compared to anti-TGFβ antibody

We investigated the binding affinity of TβR2-HFt to the recombinant human TGFβ isoforms, including TGFβ1, TGFβ2, and TGFβ3, immobilized on a solid support using ELISA. TβR2-HFt exhibited picomolar binding affinity to all three TGFβ isoforms with apparent *K*_d_ values comparable to the value observed with anti-TGFβ monoclonal antibody (mAb, clone 1D11) (**Fig. 4a** and **Table S2**). We next compared TGFβ inhibitory activity of TβR2-HFt and anti-TGFβ mAb using the reporter cell system described above where we found that IC_50_ value of TβR2-HFt (15.65 ± 0.85 pM) was markedly lower than that of anti-TGFβ mAb (210.2 ± 15.4 pM) (**Fig. 4b**). To complement this study, we assessed the ability of TβR2-HFt to deter the canonical TGFβ/SMAD signaling pathway activated by either human or mouse TGFβ1 or TGFβ2 in NIH/3T3 mouse fibroblasts and GL261-Luc2 mouse glioma cells. Western blot analysis demonstrated that all TGFβ isoforms induced phosphorylation of SMAD2 and SMAD3, indicating the activation of canonical TGFβ/SMAD signaling pathway, which was inhibited by TβR2-HFt in a dose-dependent manner (**Fig. 4c**).

**Fig. 4.**
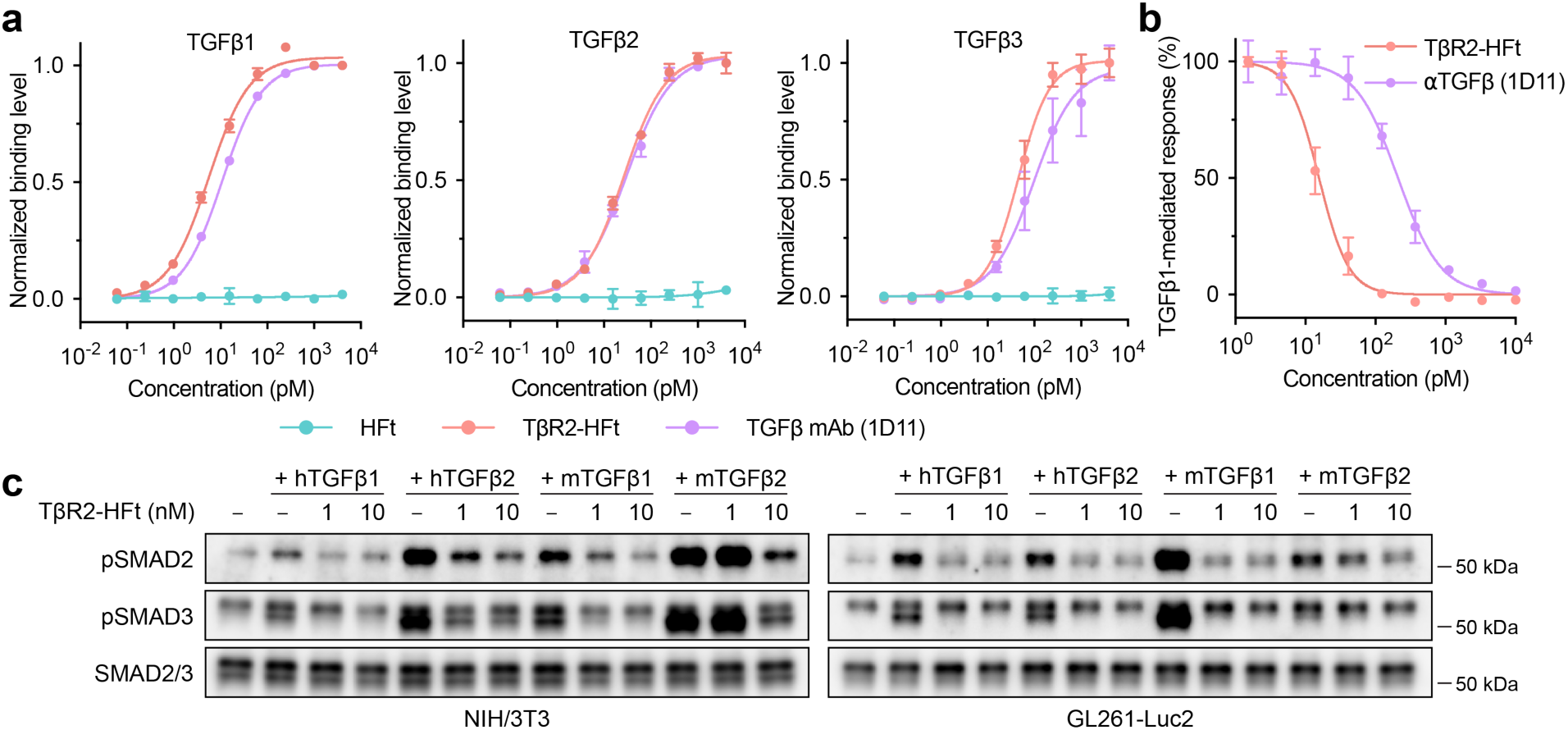
TβR2-HFt provides markedly enhanced TGFβ inhibitory activity compared to anti-TGFβ mAb while retaining high-affinity pan-TGFβ binding. **a**, Binding curves of HFt, TβR2-HFt, and anti-TGFβ mAb (αTGFβ) clone 1D11 against recombinant human TGFβ1, TGFβ2, and TGFβ3. **b,** TGFβ-inhibitory activity of TβR2-HFt and αTGFβ against recombinant human TGFβ1 in TGFβ/SMAD signaling reporter cells. **c,** Western blot analysis showing dose-dependent reduction of pSMAD2 and pSMAD3 levels (*i.e*., TGFβ-inhibitory activity) by TβR2-HFt in NIH/3T3 and GL261-Luc2 cells in presence of human (h) or mouse (m) TGFβ1/2. Data in **a** and **b** are presented as mean ± s.d.

### TβR2-HFt traverses integral BBB via receptor-mediated transcytosis

We investigated whether TβR2-HFt retains the ability of the parent HFt to bind to its receptors on brain endothelium and thus to penetrate the BBB (*21*). Pull-down assays using hCMEC/D3 human brain endothelial cell lysates confirmed that both HFt and TβR2-HFt specifically bound to human TfR1 (hTfR1) (**Fig. 5a**). It has been previously shown that HFt does not bind to mouse TfR1 (*22*) but uses mouse T-cell immunoglobulin and mucin domain-2 (mTIM-2) as a cell surface receptor (*21, 25*). We thus repeated the pull-down assay using bEnd.3 mouse brain endothelial cell lysates to confirm the robust interactions of both HFt and TβR2-HFt to mTIM-2 (**Fig. 5a**). We next evaluated the binding affinity of HFt and TβR2-HFt to recombinant hTfR1 and mTIM-2 immobilized on a solid support. HFt and TβR2-HFt showed picomolar binding affinity to both receptors with TβR2-HFt exhibiting slightly greater *K*_d_ values compared to HFt (**Fig. 5b** and **Table S3**). Reflecting their receptor binding affinity, confocal microscopy demonstrated efficient intracellular uptake of fluorescently labeled HFt and TβR2-HFt by hCMEC/D3 and bEnd.3 cells (**Fig. 5c**).

**Fig. 5.**
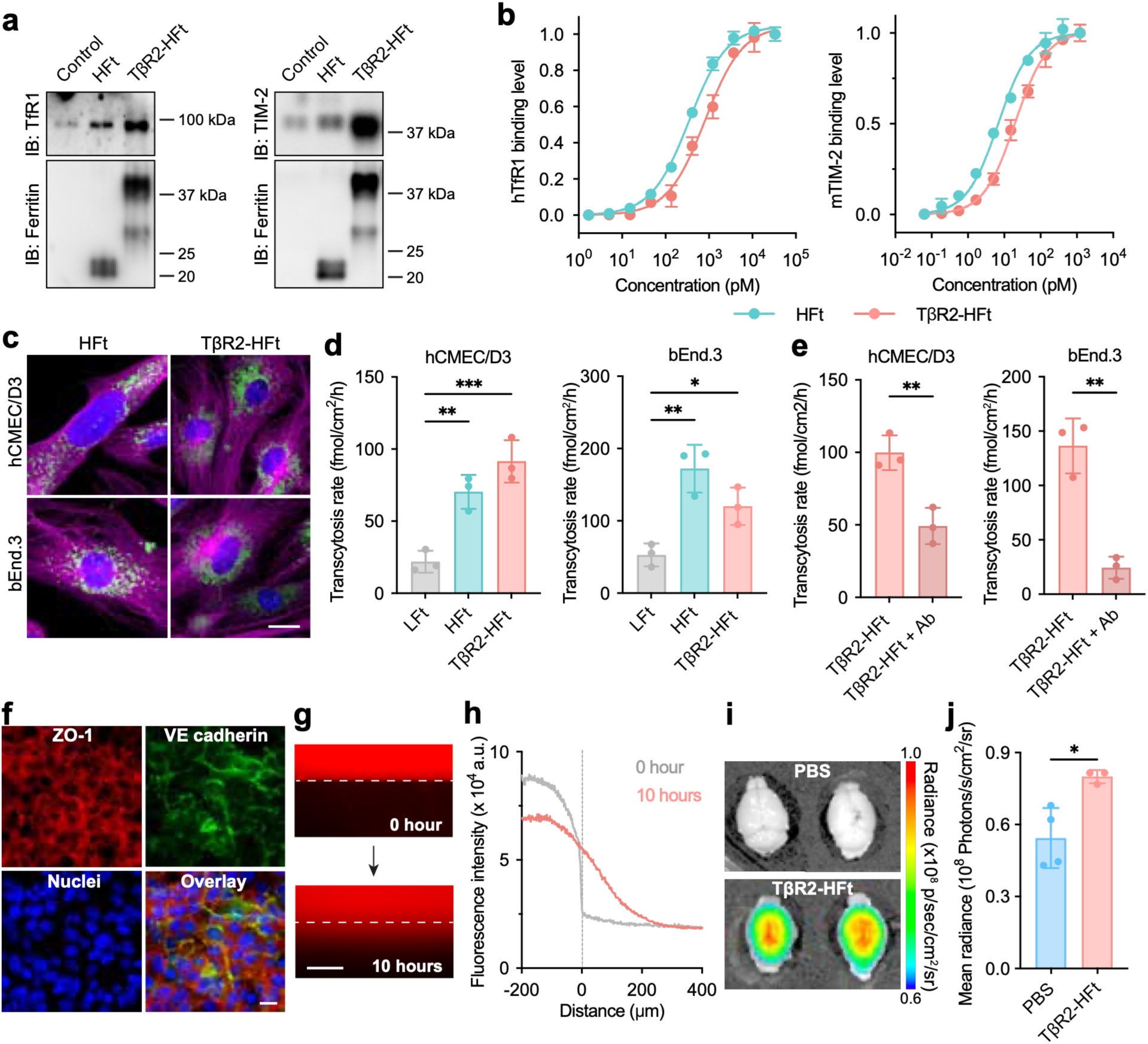
TβR2-HFt traverses integral BBB *in vitro* and *in vivo*. **a**, Western blot analysis of Ni-NTA pull-down assays showing specific interactions of HFt and TβR2-HFt with human TfR1 (hTfR1) and mouse TIM-2 (mTIM-2) in brain endothelial cell lysates. **b**, Binding curves of HFt and TβR2-HFt to recombinant hTfR1 (left) and mTIM-2 (right) immobilized on solid support. **c**, Cellular uptake of fluorescently labeled HFt and TβR2-HFt by human hCMEC/D3 (left) and mouse bEnd.3 (right) brain endothelial cells. Green, HFt/TβR2-HFt; magenta, α-tubulin; blue, nuclei. Scale bar, 20 µm. **d**, Transcytosis rates of LFt, HFt, and TβR2-HFt across *in vitro* transwell BBB models prepared with hCMEC/D3 (left) and bEnd.3 (right) cell lines. **e**, Inhibition of TβR2-HFt transcytosis across the human (left) and mouse (right) transwell BBB models by receptor blocking anti-TfR1 mAb (clone M-A712) and anti-TIM-2 mAb (clone RMT2-1), respectively. **f**, Immunostaining of junctional proteins, including ZO-1 and VE-cadherin, in a 3D microfluidic BBB chip model established with hCMEC/D3 cells and human astrocytes. Red, ZO-1; green, VE-cadherin; blue, nuclei. Scale bar, 50 µm. **g,h**, Fluorescence image showing TβR2-HFt translocation across the BBB in the 3D BBB chip model (**g**) and quantification (**h**). Scale bar, 200 µm. Red: fluorescently labeled TβR2-HFt. The white and grey dashed lines, respectively, in **g** and **h** indicate the boundary between hCMEC/D3 cells (*i.e.*, ‘blood-side’ channel) and human astrocytes in Matrigel (*i.e*., ‘brain-side’ channel) shown in Fig. S2. **i**,**j**, IVIS images showing accumulation of fluorescently labeled TβR2-HFt, following systemic administration, in healthy mouse brains with integral BBB (**i**) and quantification (**j**). Quantitative data in **b**, **d**, **e** and **j** are presented as mean ± s.d. Statistical significance was determined by one-way ANOVA with Tukey’s post hoc test for **d** or unpaired two-tailed Student’s t-test for **e** and **j**. \**p* < 0.05, \*\**p* < 0.01, \*\*\**p* < 0.001.

To test the ability of TβR2-HFt to penetrate the BBB via receptor-mediated transcytosis, we established 2D transwell BBB models using hCMEC/D3 or bEnd.3 cells (*26*). We first compared the transcytosis rates of HFt, TβR2-HFt, and a light-chain ferritin protein nanocage (LFt) as a negative control. Of note, LFt does not bind to TfR (*22*) but instead uses scavenger receptor class A member 5 as a cell surface receptor which is not expressed on brain endothelium (*27*). HFt and TβR2-HFt displayed significantly greater transcytosis rates compared to LFt in both human and mouse BBB models, and the differences between HFt and TβR2-HFt were not significant (**Fig. 5d**). We also found that pre-treatment with blocking antibodies against hTfR1 or mTIM-2 significantly reduced the transport of TβR2-HFt across the human or mouse brain endothelial monolayers, respectively, indicating that transcytosis took place in a receptor-dependent manner (**Fig. 5e**). To complement the 2D transwell BBB model study, we investigated the permeability of TβR2-HFt in a 3D microfluidic BBB chip model, which better recapitulates unperturbed BBB. The model is composed of a ‘blood-side’ channel lined with hCMEC/D3 cells and a ‘brain-side’ compartment filled with human astrocytes embedded in Matrigel, separated by a well-defined boundary (**Fig. S2**). A tightly sealed BBB monolayer was confirmed by immunofluorescence showing uniform expression of tight junction molecules, including zonula occludens-1 and vascular endothelial cadherin (**Fig. 5f**). We found that fluorescently labeled TβR2-HFt added to the blood-side channel gradually traversed the endothelial barrier in the model to reach the brain-side compartment over time (**Figs. 5g, h**).

We next tested the ability of TβR2-HFt to penetrate the BBB *in vivo*. An orthotopic mouse glioma model was established by intracranial inoculation with GL261-Luc2 cells, and individual tumor-bearing mice were treated with fluorescently labeled TβR2-HFt via systemic administration. Live-animal fluorescence imaging revealed accumulation of the nanocages in the brain tumor characterized by leaky BBB (**Fig. S3a**). In parallel, we confirmed using *ex vivo* IVIS imaging of the harvested brain tissues that fluorescently labeled TβR2-HFt, following systemic administration, accumulated in healthy mouse brains with integral BBB (**Figs. 5i, j**).

### TβR2-HFt efficiently penetrates 3D multicellular tumor spheroids and binds to human and mouse glioma cells *in vitro*

We have previously demonstrated that HFt is capable of penetrating non-brain tumor tissues *in vitro* and *in vivo* and have established evidence that the penetration is likely mediated by its receptors present on tumor cells (*25, 28*). We thus tested whether TβR2-HFt retained the ability of the parent HFt to penetrate 3D multicellular spheroids. Fluorescently labeled TβR2-HFt exhibited widespread distribution throughout the spheroid based on GL261-Luc2 cells to a similar extent as HFt (**Fig. S3b**). We next investigated the ability of TβR2-HFt to bind to human (HSR-GBM1) and mouse (GL261-Luc2 and CT-2A) glioma cell surfaces. To do so, cells were incubated with fluorescently labeled TβR2-HFt at 4 °C to minimize internalization. Subsequent flow cytometry analysis revealed surface binding of the nanocages with all three cell types, as evidenced by the cell-associated fluorescence (**Fig. S4**).

### Systemically administered TβR2-HFt inhibits canonical TGFβ/SMAD signaling pathway and mitigates immunosuppressive TME via modulation of both stromal and immune components in an orthotopic mouse glioma model

To investigate the ability of TβR2-HFt to modulate immunosuppressive TME in GBM, we established an orthotopic glioma model by intracranial inoculation of GL261-Luc2 cells into the right hemisphere. In agreement with the human data (Fig. 2), *Tgfb1*, *Tgfb2*, and *Tgfbr2* mRNA transcript levels were greater in tumor-bearing hemispheres than in contralateral tumor-free hemispheres while the difference was not statistically significant for *Tgfb2* (**Fig. S5**). In parallel, we conducted a dose-escalation safety study using healthy mice to confirm that systemic TβR2-HFt dose of 10 mg/kg did not induce body weight loss upon repeated treatments (**Fig. S6**). We thus established the model again and treated individual tumor-bearing mice with PBS or TβR2-HFt at a dose of 10 mg/kg via intravenous injection every 5 days up to four doses. Immunohistochemical analysis of tumor-bearing brain tissues revealed that TβR2-HFt significantly reduced the phosphorylated SMAD (pSMAD)-positive area in comparison to PBS-treated tumor-bearing brain tissues (**Figs. 6a, b**), in agreement with our *in vitro* observations (**Figs. 3e, 4b, c**).

**Fig. 6.**
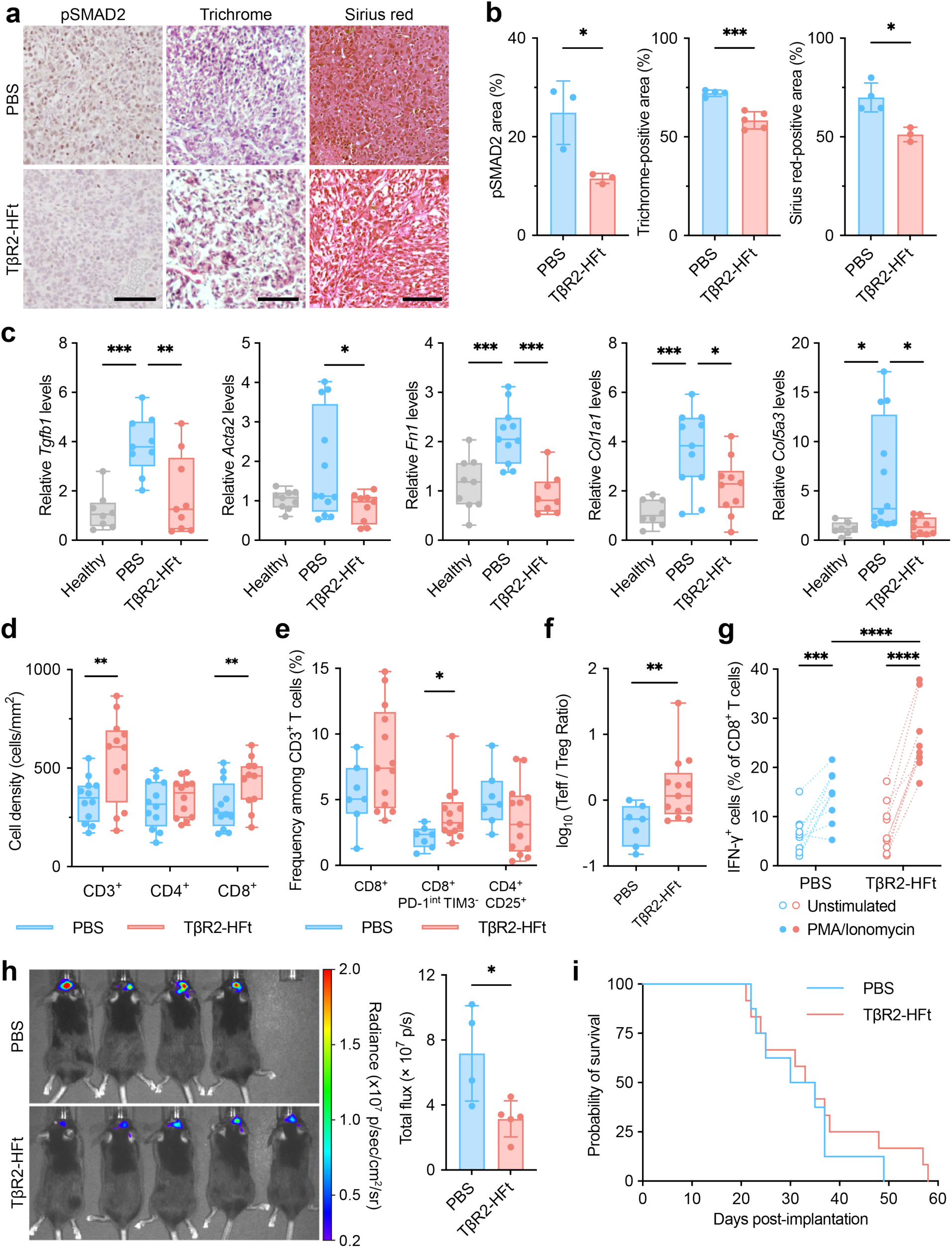
Systemically administered TβR2-HFt mitigates immunosuppressive TME in an orthotopic mouse glioma model. **a,b,** Representative pSMAD2 immunohistochemistry (left) and Masson’s trichrome (middle) and Sirius red (right) staining of brain tumor sections from mice treated with PBS or TβR2-HFt (**a**) and the quantification of the stained areas (**b**). Scale bar, 100 μm. **c**, Relative mRNA transcript levels of *Tgfb1*, *Col1a1*, *Col5a3*, *Fn1*, and *Acta2* in healthy brains and tumor-bearing brains from mice treated with PBS or TβR2-HFt. **d**, Density of CD3^+^, CD4^+^, and CD8^+^ tumor-infiltrating lymphocytes (cells/mm²). **e**, Flow cytometry analysis showing the fractions of the overall CD8^+^ T (CD3^+^ CD8^+^), effector CD8^+^ T (CD3^+^ CD8^+^ PD-1^int^ TIM3^-^), and regulatory T (Treg, CD3^+^ CD4^+^ CD25^+^) cells in tumors. **f**, A logarithm of the intratumoral ratio of effector CD8+ T cells to Treg cells. **g**, Flow cytometry analysis showing the fraction of IFN-γ-producing CD8^+^ T cells isolated from tumors and subsequently stimulated *ex vivo* with PMA/ionomycin. **h**, *In vivo* bioluminescence images showing GL261-Luc2 cells in mouse brains (left) and the quantification of total flux (right). **i**, Kaplan-Meier survival analysis. Data in **b** and **h** are presented as mean ± s.d. Box-and-whisker plots in **c**-**f** show the median and interquartile range, with the whiskers indicating the minimum and maximum values. Statistical significance was determined by unpaired two-tailed Student’s t-test for **b**, **d**-**f** and **h**, one-way ANOVA with Tukey’s post hoc test for **c**, two-way repeated measures ANOVA with Sidak’s multiple comparisons test for **g**, and the log-rank test for **i**. \**p* < 0.05, \*\**p* < 0.01, \*\*\**p* < 0.001, \*\*\*\**p* < 0.0001.

Upon the confirmation of *in vivo* TGFβ inhibitory activity, we evaluated whether systemic TβR2-HFt treatment altered the stromal component of immunosuppressive TME of the model. We found that TβR2-HFt significantly reduced Masson’s trichrome- and Sirius red-stained collagen-positive areas compared to the PBS-treated control (**Figs. 6a, b**). In addition, tumor-bearing brain tissues from TβR2-HFt-treated mice exhibited significant reductions of the levels of *Tgfb1*, *Acta2*, *Fn1*, *Col1a1*, and *Col5a3* mRNA transcripts compared to PBS-treated tumor-bearing brain tissues, towards the levels observed in healthy brain tissues (**Fig. 6c**). We next examined the impact of systemic TβR2-HFt treatment on the immune cell profile in immunosuppressive TME of the model. Quantitative analysis of immunostained tumor tissue sections showed increased densities of CD3^+^ and CD8^+^ T cells in TβR2-HFt-treated tumor tissues compared to PBS-treated tumor tissues (**Fig. 6d**). We further analyzed the distinct T-cell populations in TβR2-HFt-treated versus PBS-treated tumor-bearing brain tissues using flow cytometry analysis. Notably, intratumoral effector T cell (CD8^+^PD-1^int^ TIM3^−^) fraction was significantly increased by TβR2-HFt treatment, which led to a concomitant increase in the fractional ratio of effector T cells over Treg cells (CD4^+^CD25^+^) (**Figs. 6e, f**). Effector T cells were then subjected to an *ex vivo* PMA/ionomycin restimulation study to show a greater frequency of interferon-γ (IFN-γ)-producing cells when TβR2-HFt treatment was given (**Fig. 6g**). Using a separate but identically treated animal cohort, we monitored the tumor burden and survival. While *in vivo* bioluminescent imaging demonstrated moderate but significant reduction of tumor burden by TβR2-HFt treatment, the treatment alone failed to confer survival extension of tumor-bearing mice (**Figs. 6h, i**), despite its ability to extensively mitigate immunosuppressive TME (**Figs. 6c-g**).

### TβR2-HFt downregulates stromal fibrosis markers in human glioma spheroids and astrocytes

To establish the relevance of the observed *in vivo* TME modulation to human GBM, we assessed whether TβR2-HFt would mitigate TGFβ-induced stromal activation in human glioma cells and astrocytes. Of note, it is now increasingly appreciated that astrocytes are activated in the TME of GBM to promote tumor growth and invasion (*29*), and activated astrocytes are shown to upregulate stromal fibrosis-associated proteins (*30, 31*). We found that TβR2-HFt suppressed TGFβ1-induced transcription of *TGFB1*, *CTGF*, and *SERPINE1* in HSR-GBM1 spheroids (**Fig. S7a**). Likewise, the treatment blunted TGFβ1-induced upregulation of *TGFB1*, *CTGF*, *SERPINE1*, *COL1A1*, and *FN1* in H9 human embryonic stem cell (hESC)-derived human astrocytes (**Fig. S7b**).

### Systemically administered TβR2-HFt rescues the anti-cancer therapeutic efficacy of a T-cell ICI in multiple orthotopic mouse glioma models

We investigated whether systemic TβR2-HFt-mediated modulation of TME would enhance the therapeutic efficacy of anti-programmed cell death protein 1 (PD-1) mAb in multiple orthotopic mouse glioma models. We first tested the combination therapy in a model based on GL261-Luc2 cells. Mice with comparable tumor burdens were randomly assigned to four groups and treated with PBS, anti-PD-1 mAb (clone RMP1-14, 10 mg/kg hereafter), a combination of anti-TGFβ mAb (clone 1D11, 10 mg/kg) and anti-PD-1 mAb, or a combination of TβR2-HFt (10 mg/kg) and anti-PD-1 mAb every 5 days and monitored for survival. Presumably reflecting the immunosuppressive nature of the model (Fig. 6), anti-PD-1 mAb monotherapy failed to significantly extend the survival of the model (**Fig. 7a** and **Table S4**). However, while an addition of anti-TGFβ mAb was unable to prolong the survival, the combined treatment with TβR2-HFt and anti-PD-1 mAb significantly extended the survival of GL261-Luc2 tumor-bearing mice (**Fig. 7a** and **Table S4**). We next evaluated the therapeutic efficacy of our therapeutic strategy in an additional orthotopic mouse glioma model prepared via intracranial inoculation of CT-2A cells. A CT-2A model, due to its more immunosuppressive and less immunogenic nature, has been shown to exhibit a poorer response against anti-PD-1 mAb compared to a GL261 model (*32*). After confirming tumor implantation via magnetic resonance imaging (MRI), the animals with comparable tumor burdens were assigned to three groups and treated with PBS, anti-PD-1 mAb, or a combination of TβR2-HFt and anti-PD-1 mAb every two days up to three overall doses. Similar to our observation with a GL261 model, anti-PD-1 mAb monotherapy was unable to significantly prolong the survival, but a statistically significant survival extension was achieved by the combination of TβR2-HFt and anti-PD-1 mAb, yielding a fraction of long-term survival mice (>120 days, 2 out of 9 mice) (**Fig. 7b** and **Table S4**).

**Fig. 7.**
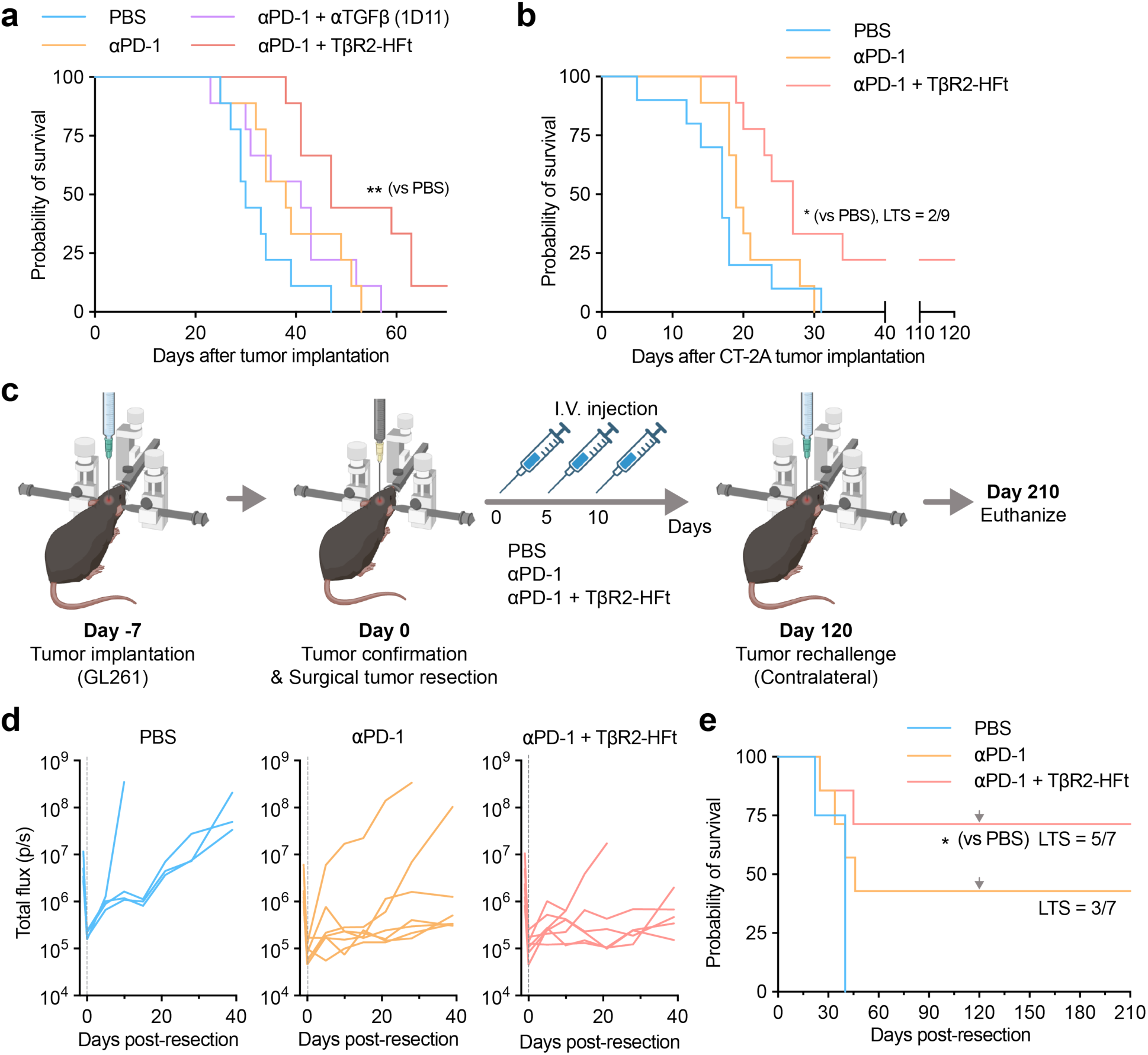
Systemically administered TβR2-HFt potentiates anti-PD-1 immunotherapy in orthotopic primary glioma models and a post-resection glioma model with tumor rechallenge. **a,** Kaplan-Meier survival analysis of orthotopic GL261-Luc2 tumor-bearing mice treated with PBS, anti-PD-1 mAb (αPD-1), αPD-1 plus αTGFβ clone 1D11, or αPD-1 plus TβR2-HFt (n = 9 per group). **b,** Kaplan-Meier survival analysis of orthotopic CT-2A tumor-bearing mice treated with PBS, αPD-1, or αPD-1 plus TβR2-HFt (PBS, n = 10; other groups, n = 9). **c,** Experimental timeline showing initial tumor inoculation, surgical tumor resection, post-resection treatments, and contralateral orthotopic tumor rechallenge. Days are shown relative to the day of surgical resection. **d,** Longitudinal bioluminescence imaging of recurrent GL261-Luc2 tumors following surgical resection. **e,** Kaplan-Meier survival analysis of orthotopic GL261-Luc2 tumor-bearing mice that received surgical tumor resection and subsequent treatments with PBS, αPD-1, or αPD-1 plus TβR2-HFt (PBS, n = 4; αPD-1, n = 7; αPD-1 plus TβR2-HFt, n = 7). Gray arrows indicate contralateral orthotopic tumor rechallenge of long-term survival mice on day 120 after resection. Statistical significance was determined by the log-rank test with Bonferroni correction for multiple comparisons.

As introduced earlier, most of the GBM patients undergo surgical tumor resection but near-inevitably encounter lethal tumor recurrence (*1, 33*). To reflect this clinical scenario, we surgically removed tumors from orthotopic GL261-Luc2 tumor-bearing mice and treated them with PBS, anti-PD-1 mAb, or a combination of TβR2-HFt and anti-PD-1 mAb every 5 days up to three overall doses (**Fig. 7c**). Although anti-PD-1 mAb monotherapy exhibited a median survival comparable to PBS-treated group, roughly half of the mice (3 out of 7 mice) survived the tumor with enhanced control over tumor growth (**Figs. 7d, e** and **Table S4**). Notably, the combined treatment with TβR2-HFt and anti-PD-1 mAb enabled a statistically significant survival extension, yielding a majority of long-term survival mice (5 out of 7 mice) demonstrating remarkable tumor suppression (**Figs. 7d, e** and **Table S4**). The long-term survival mice in both groups were then rechallenged with contralateral orthotopic GL261-Luc2 inoculation to assess whether the treatments endowed the animals with the ability to control the tumor in a durable manner. All rechallenged mice in both treatment groups remained alive for at least 90 days post-rechallenge without additional intervention (**Figs. 7e** and **S8**). In contrast, all age-matched control mice that received identical tumor inoculation succumbed to death (**Fig. S8**).

## Discussion

We here demonstrate that our newly engineered TβR2-HFt, following systemic administration, reverses the immunosuppressive TME to potentiate otherwise inefficacious anti-PD-1 mAb, an ICI analogous to a clinical immunotherapy used against multiple non-brain tumors (*2*). Using complementary experimental models, we validated our hypothesis that TβR2-HFt would address key delivery challenges and promote multivalent TGFβ trapping to remodel both stromal and immune components of immunosuppressive TME in GBM. Notably, TβR2-HFt near-fully retained the ability to engage with inherent molecular targets of both the delivery (*i.e*., FHC) and the therapeutic (*i.e*., TGFβR2ecd) elements, thereby enabling the remarkable *in situ* immunomodulatory outcomes. Our findings underscore the versatility of the platform, which could be potentially applied to other immunomodulatory targets and/or paired with various immunotherapeutic modalities currently used in clinic or under clinical investigation.

It appears that the unique ability of TβR2-HFt to traverse the BBB is critical to achieving effective intratumoral TGFβ inhibition and restoration of ICI efficacy. Our *in vitro* and *in vivo* studies collectively show that TβR2-HFt is capable of penetrating the BBB via receptor-mediated transcytosis regardless of its integrity and across species. The importance of such ability is underscored by our observation that an antibody-based TGFβ inhibitor (*i.e*., anti-TGFβ mAb) is unable to rescue the ICI efficacy despite tumor-induced BBB disruption in GBM (*19*). The finding is well aligned with the previous reports demonstrating that circulating antibodies cannot readily traverse the intact BBB (*34, 35*). While antibody-based T-cell ICIs, such as anti-PD-1 mAb, gain access to the brain tumor primarily via circulating T cells (*36*), antibody therapies that directly target cellular or non-cellular components within brain tumors, such as anti-TGFβ mAb and cancer cell-targeting antibodies, require BBB penetration following systemic administration (*34, 35*). As introduced above, BBB disruption is spatially heterogeneous in GBM, and invasive tumor cells infiltrate into the surrounding normal brain parenchyma where the barrier function of BBB remains largely integral (*19*). Given that those infiltrating tumor cells near-inevitably reinitiate lethal recurrent GBM, the ability to reach and tackle those regions is a prerequisite to realizing durable tumor control and potential cure (*37, 38*).

Once beyond the BBB, the ability of TβR2-HFt to penetrate the brain tumor tissue and bind to brain tumor cells likely contributed to widespread and efficacious intratumoral TGFβ inhibition. TβR2-HFt exhibited uniform coverage in 3D multicellular spheroids established with GL261-Luc2 mouse glioma cells, to an extent virtually identical to its parent HFt. We have previously demonstrated that HFt is capable of efficiently penetrating non-brain tumor tissue *in vivo* and 3D multicellular spheroids *in vitro*, including hypoxic tumor regions, in a receptor-dependent manner (*25, 28*). Of note, the abnormal vasculature in GBM creates hypo-perfused tumor areas, including hypoxic regions (*39*), that are not readily accessible by systemically administered therapeutics (*40*). On the other hand, TGFβ is upregulated in human brain tumor cells to promote immunosuppression under hypoxia (*41, 42*), which highlights the importance of achieving extensive TGFβ inhibition throughout the tumor tissue. In addition to the tumor-penetrating ability, our flow cytometry analysis revealed that TβR2-HFt efficiently binds to both human (*i.e*., HSR-GBM1) and mouse (*i.e*., GL261-Luc2 and CT-2A) glioma cells. We speculate that the ability of TβR2-HFt to anchor on brain tumor cells enhances its intratumoral retention while endowing a “gatekeeping” function to trap incoming and outgoing TGFβ, thereby enabling prolonged and therapeutically viable TGFβ inhibition.

The remarkable TGFβ inhibition was also achieved by multivalent TGFβR2ecd display on the nanocage platform. As introduced earlier, HFt is formed by self-assembly of 24 protein subunits (*23*), and thus up to 24 TGFβ trapping domains can be theoretically displayed on each TβR2-HFt surface, in comparison to divalent anti-TGFβ mAb. Our *in vitro* TGFβ/SMAD reporter assay revealed that the IC_50_ value of TβR2-HFt was over 13-fold lower than that of anti-TGFβ mAb, suggesting that the multivalent effect could be beyond additive. Such multivalent effect was enabled by careful fusion protein design where the nanocage based on fusion protein variant comprising TGFβR2ecd fused to the C-terminus of FHC exhibited excellent TGFβ binding and inhibitory activities, but those with an opposite domain order failed to do so. Likewise, TGFβR2ecd fused to the C-termini of other proteins, including human serum albumin, has been shown to preserve its biological activities (*14, 43*). The C-terminus of FHC orients towards the inner cavity of the nanocage, but a peptide with ≥ 90 amino acids fused to the C-terminus has been reported to be extruded outside due to the limited cavity volume to accommodate the peptides from 24 protein subunits (*44*). Thus, 137-amino acid TGFβR2ecd was expected to be displayed on the nanocage surface, which was validated by the greater hydrodynamic diameter of TβR2-HFt compared to HFt. In agreement with our observation, a protein or a peptide C-terminally fused to FHC has been shown to be displayed on the nanocage surface to exert a respective biological function (*45, 46*). We also confirmed that the C-terminal fusion of TGFβR2ecd marginally reduced the binding affinity of HFt to hTfR1. This minimal interference is likely due to the presence of the hTfR1 contact regions near the surface exposed N-terminal FHC (*47*), which may not be masked significantly by the C-terminally fused TGFβR2ecd.

TβR2-HFt, by delivery enhancement and multivalent TGFβ inhibition, concurrently reversed tumor-promoting stromal and immune events manifested in immunosuppressive TME of GBM. TβR2-HFt de-densified the collagen-rich tumor stromal ECM and reduced fibrosis-associated mRNA transcript levels in orthotopic gliomas. This comprehensive stromal modulation was accompanied by increased intratumoral CD3^+^ and CD8^+^ T-cell densities, in agreement with previous studies demonstrating that TGFβ inhibition promoted immune cell infiltration into tumor tissues of multiple non-brain tumor models (*10–12*). In parallel, TGFβ-induced pro-fibrotic factors were downregulated in HSR-GBM1 human glioma spheroids and human embryonic stem cell-derived astrocytes by TβR2-HFt *in vitro*. The findings underlie its translational potential to mitigate the stromal barrier by acting on both malignant and non-malignant cellular compartments within TME of human GBM. TβR2-HFt also altered the intratumoral T-cell profile to enrich tumor-killing effector T cells and increase the effector-T-to-Treg-cell ratio *in vivo*. Of note, the canonical TGFβ/SMAD signaling pathway has been shown to promote T-cell exhaustion (*48, 49*) and Treg-cell induction (*50, 51*) in numerous *in vitro* and *in vivo* non-brain tumor model studies, suggesting that the favorable T-cell modulation observed in this study is likely attributed to the robust ability of TβR2-HFt to deactivate the pathway.

We demonstrate here that TβR2-HFt consistently rescues the therapeutic efficacy of an anti-PD-1 mAb across cell types (*i.e.*, GL261 and CT-2A) and clinical stages (*i.e*., pre- and post-resection) of orthotopic mouse gliomas. The combination therapy exhibited a more pronounced therapeutic outcome in a primary CT-2A model compared to a stage-matched GL261 model, despite the greater immunosuppressive and less immunogenic nature of the former (*32*). The observation is presumably due to a more frequent dosing schedule that we have implemented on the CT-2A model accounting for its enhanced aggressiveness compared to GL261 models (*32*). Intratumoral TGFβ level and its impact on immunosuppressive TME are expected to vary across glioma types, which may have additionally contributed to the outcome. Remarkably, a majority (>70%) of GL261 tumor-bearing mice that received surgical resection and subsequent combination therapy resisted tumor recurrence and rechallenge, suggesting that durable glioma-specific immune memory was established. Albeit to a lesser extent, a fraction of the tumor-resected mice treated with anti-PD-1 mAb monotherapy also showed such resistance, in contrast to its negligible benefit observed among recurrent GBM patients who received surgical tumor resection and standard-of-care radio- and/or chemotherapy (*7, 52*). Thus, while this study establishes solid proof-of-concept efficacy and mechanism, validation of our strategy in models better emulating human GBM and clinical scenario is warranted. Of note, we showed that TβR2-HFt treatment increased IFN-γ-producing T cells in TME of a mouse glioma model. This finding well aligns with a recent clinical study demonstrating the enrichment of the IFN-γ signaling pathway in recurrent GBM patients that responded to an investigational dual-ICI therapy (*53*), providing a rationale to further develop our immunomodulatory strategy for enhanced immunotherapy of GBM.

## Materials and Methods

### Preparation and characterization of recombinant proteins and nanocages

The fusion protein variants of FHC and TGFβR2ecd (Thr23-Asp184, NCBI accession no. NP_001020018) with different domain orders and linker types, including a flexible (GGGGS)_4_ or a rigid A(EAAAK)_4_ALEA(EAAAK)_4_A linker, were constructed. Plasmid DNA vectors encoding fusion protein variants were synthesized by ATUM. Briefly, DNA sequences for the fusion protein variants were codon-optimized for bacterial expression using GeneGPS^®^ and cloned into the pD861-NH plasmid DNA vector with an N-terminal 6xHis tag. A TEV protease-cleavable sequence was inserted between the 6xHis tag and the fusion protein construct to allow removal of the 6xHis tag, if needed (**Table S1**). Each vector was transformed into competent BL21(DE3) *E. coli* cells (Thermo Fisher Scientific, Cat. EC0114) by heat shock, and transformants were selected for kanamycin resistance. Structural models of the fusion protein subunit were generated using the AlphaFold server (*54*). The multimeric nanocage structure was generated by ChimeraX 1.10.1 (*55*).

Cells were grown in LB broth containing kanamycin (50 µg/mL) to an OD₆₀₀ of 0.6-0.8, and protein expression was induced with 1 mM L-rhamnose for 5 h at 37 °C. Harvested cells were pelleted by centrifugation (5,000 × g, 10 min). After washing with cold PBS, the cells were lysed using B-PER™ Complete reagent (Thermo Fisher Scientific). Insoluble inclusion bodies were harvested (16,000 × g, 15 min), washed with TBS containing 1% Triton X-100, and solubilized overnight at 4 °C in denaturing buffer (50 mM Tris-HCl, 8 M urea, 5 mM DTT, pH 8.0). Lysates (adjusted to 5 mg/mL) were diluted 10-fold into refolding buffer (50 mM Tris-HCl, 0.5 M L-arginine, 2 mM GSH, 0.2 mM GSSG, pH 8.0) and stirred for 72 h at 4 °C. Refolded proteins were purified via Ni-NTA affinity chromatography (Qiagen) with 250 mM imidazole elution, dialyzed against PBS, and purified by preparative SEC on a Superose 6 Increase 10/300 GL column (Cytiva). Protein expression and final purity were assessed using reducing SDS-PAGE with Coomassie blue staining. The hydrodynamic diameters of nanocages formed by self-assembly of the recombinant proteins were determined by DLS using a Zetasizer Nano ZS90 (Malvern Instruments). Analytical SEC was performed on a Shimadzu LC-2030C system using a MabPac SEC-1 column (4 × 300 mm; Thermo Fisher Scientific) in mobile phase (50 mM sodium phosphate, 300 mM NaCl, pH 6.8) at 1 mL/min. Unless otherwise indicated, recombinant proteins were used with the N-terminal 6xHis tag retained.

### Analysis of public database for the mRNA transcript levels in human glioma

Log_2_-transformed DESeq2-normalized RNA-seq data for TGFB1, TGFB2, TGFB3, TGFBR1, and TGFBR2 were obtained from the UCSC Toil recompute hub via the UCSC Xena browser (*56*). Primary TCGA-GBM tissue samples (*n* = 153) were compared with GTEx normal brain tissues (*n* = 1,142) and TCGA-LGG tissue samples; recurrent GBM (*n* = 13) and adjacent normal tissues (*n* = 5) were excluded due to limited sample sizes.

Correlations between mRNA transcript levels of TGFβ isoforms and those of TGFβ receptors and stromal fibrosis and immune suppression markers in TCGA-GBM and CGGA-GBM cohorts were analyzed using the GlioVis portal (*57*). Analyses were performed using RNA-seq data with the following inclusion criteria: adult patient, the presence of GBM histology data, and all subtypes. The TGFβ score was defined using *TGFB1*, *TGFB2*, and *TGFB3* mRNA transcript levels on average. Stromal fibrosis (*FN1*, *COL1A1*, *COL5A3*, and *ACTA2*) and immune suppression (*CD274*, *HAVCR2*, *FOXP3*, and *IL2RA*) scores were calculated from the respective mRNA transcript levels on average. Pairwise correlations were assessed using Pearson correlation coefficients. For expanded analyses shown in the supplementary figures, additional stromal fibrosis (*SERPINE1*, *CTGF*, *COL6A3*, and *FAP*) and the immune suppression (*CD163*) markers were included.

### *In vitro* TGFβ/SMAD activity assay

HEK293T cells (ATCC) were reverse-transfected with pGL4.48 (Promega) using FuGENE HD (3 µL per 1 µg DNA; Promega). Transfected cells were seeded in 96-well plates (1 × 10^4^ cells per well) and cultured overnight at 37 °C. Human or mouse TGFβ1 (5 ng/mL; PeproTech, Cat. 100-21) and serial dilutions of test compounds were added to the cells, followed by overnight incubation at 37 °C. Luminescence was measured on a microplate reader following a 5-min incubation with Steady-Glo reagent (Promega). IC_50_ values were determined by nonlinear regression fitting to a four-parameter logistic model in GraphPad Prism. Nanocage concentrations were calculated based on the molecular weight of the fully assembled 24-mer cage.

### *In vitro* binding assays using ELISA

For TGFβ binding assays, high-binding 96-well plates (Corning) were coated overnight at 4 °C with human TGFβ1, TGFβ2, or TGFβ3 (0.5 µg/mL; PeproTech). For receptor-binding assays, MaxiSorp 96-well plates (Thermo Fisher Scientific) were coated overnight at 4 °C with goat anti-human IgG Fcγ antibody (10 µg/mL; Jackson ImmunoResearch, Cat. 109-005-008) and subsequently incubated with recombinant hTfR1-Fc (2 µg/mL; Sino Biological, Cat. 11020-H01H) or mTIM-2-Fc (2 µg/mL; BioVision, Cat. P1355) for 2 h at room temperature. All plates were washed with PBST (PBS with 0.05% Tween 20) and blocked with 3% BSA in PBST for 1 h. Serial dilutions of test compounds were added to the wells. Bound TβR2-HFt was detected using rabbit anti-ferritin antibody (1:1000; Abcam, Cat. ab75973), followed by HRP-conjugated anti-rabbit IgG (1:10,000; Abcam, Cat. ab6721); bound anti-TGFβ mAb (clone 1D11.16.8; BioXCell, Cat. BE0057) was detected with HRP-conjugated anti-mouse IgG (1:10,000; Abcam, Cat. ab97023); biotinylated nanocages (labeled using EZ-Link NHS-PEG4 Biotinylation Kit, Thermo Fisher Scientific) were detected with streptavidin-HRP (1:4000; Thermo Fisher Scientific, Cat. 21130). Reactions were developed with TMB substrate (Thermo Fisher Scientific), quenched with 1 N sulfuric acid, and measured at 450 nm. Apparent *K*_d_ values were calculated by nonlinear regression fitting in GraphPad Prism based on assembled 24-mer nanocage molar concentrations.

### Western blot analysis of SMAD phosphorylation

NIH/3T3 mouse fibroblasts and GL261-Luc2 mouse glioma cells were serum-starved for 6 h in serum-free DMEM and treated with mouse TGFβ1 or TGFβ2 (5 ng/mL; Cell Signaling Technology, Cat. 5232LC; R&D Systems, Cat. 7346-B2-005) or human TGFβ1 or TGFβ2 (5 ng/mL; PeproTech, Cat. 100-21 and 100-35B), with or without TβR2-HFt (0.1, 1, or 10 nM). After incubation for 15 min at 37 °C, cells were washed with cold PBS and lysed using RIPA buffer supplemented with protease and phosphatase inhibitor cocktails (Thermo Fisher Scientific). The samples were resolved by SDS-PAGE and transferred to nitrocellulose membranes. The membranes were probed with primary antibodies against pSMAD2 (1:1000; Cell Signaling Technology, Cat. 3108), pSMAD3 (1:1000; Abcam, Cat. ab52903), and SMAD2/3 (1:1000; Cell Signaling Technology, Cat. 8685). After incubation with species-matched HRP-conjugated secondary antibodies, protein bands were developed using Bio-Rad ECL reagents and imaged with a Bio-Rad ChemiDoc imaging system.

### Pull-down assay

The binding of TβR2-HFt to hTfR1 and mTIM-2 was assessed by pull-down assay using hCMEC/D3 (Merck, Cat. SCC066) and bEnd.3 (ATCC, Cat. CRL-2299) cell lysates, respectively. Ni-NTA magnetic beads (Thermo Fisher Scientific) were coated with His-tagged HFt or TβR2-HFt, and uncoated beads served as controls. Coated beads were incubated for 30 min with respective cell lysates prepared using Pierce IP lysis buffer (Thermo Fisher Scientific). Beads were washed with buffer (50 mM phosphate, 300 mM NaCl, 10 mM imidazole, 0.1% Tween 20, pH 7.4) and eluted with PBS containing 250 mM imidazole for 30 min. Eluted supernatants were analyzed by western blotting. The membranes were probed with primary antibodies against ferritin (Abcam, Cat. ab75973), hTfR1 (Abcam, Cat. ab214039), and mTIM-2 (Abcam, Cat. ab86480). Protein bands were detected as described above.

### Confocal microscopy for cellular uptake analysis

HFt and TβR2-HFt were fluorescently labeled using Alexa Fluor 555 (AF555) NHS ester (Thermo Fisher Scientific). hCMEC/D3 and bEnd.3 cells were seeded on coverslips precoated with rat-tail type I collagen (Thermo Fisher Scientific) and incubated with AF555-labeled HFt or TβR2-HFt (40 nM) for 4 h at 37 °C. Fixed cells were stained with anti-α-tubulin antibody (1:2000; Cell Signaling Technology, Cat. 3873) and fluorophore-conjugated secondary antibody. Nuclei were counterstained with DAPI. Coverslips were mounted using ProLong Glass Antifade Mountant (Thermo Fisher Scientific) and imaged on a confocal microscope (LSM 710, Carl Zeiss).

### *In vitro* 2D transwell model study

Transwell membranes were precoated with rat tail collagen I and seeded with hCMEC/D3 or bEnd.3 cells (5 × 10^4^ cells/cm^2^). Cells were cultured in the upper chamber for 5-7 days at 37 °C until transendothelial electrical resistance exceeded 30 Ω·cm^2^. Test compounds (LFt, HFt, or TβR2-HFt) were added to the upper chamber at 10 nM, and samples were collected from the lower chamber at the indicated time points. For receptor-blocking studies, cells were treated with a 10-fold molar excess of mouse anti-human CD71 mAb (clone M-A712; BD Pharmingen, Cat. 555534) or rabbit anti-TIM-2 antibody (Abcam, Cat. ab86480) in the human or mouse BBB model, respectively, before addition of TβR2-HFt. Transported nanocages were quantified by sandwich ELISA. Samples and standards were captured on nickel-coated 96-well plates (Thermo Fisher Scientific) overnight at 4 °C. Plates were washed with PBST and probed with rabbit anti-ferritin antibody (1:1000; Abcam, Cat. ab75973) overnight at 4 °C. Wells were then incubated with HRP-conjugated secondary antibody for 2 h at room temperature, developed with TMB substrate, quenched with 1 N sulfuric acid, and read at 450 nm. Transcytosis rates were calculated by normalizing the recovered protein amount to membrane surface area and incubation time (fmol/cm²/h). Assays were performed using *n* = 3 independent transwell inserts per group.

### *In vitro* 3D microfluidic BBB chip model study

PDMS microfluidic devices were fabricated using 3D-printed master molds as previously described (*58*), bonded to glass slides by oxygen plasma treatment, and UV-sterilized for 3 min. Human astrocytes (0.4 × 10^4^ cells/µL; ScienCell, Cat. 1800) mixed with Matrigel (Corning) at a 1:2 volume ratio (20 μL) were injected into the brain-side channel on ice and gelled at 37 °C for 24 h, followed by perfusion of the blood-side channel with 10% Matrigel medium for 24 h. After 3 days of astrocyte culture, hCMEC/D3 cells (100 μL at 2 × 10^6^ cells/mL; Merck) were seeded into the blood-side channel and tilted at 60° overnight to facilitate attachment. The medium was refreshed daily with complete EndoGRO-MV containing 1.5 μM CHIR99021 for 3 days to establish a confluent endothelial monolayer. To verify barrier integrity, separate chips were fixed with 4% paraformaldehyde, permeabilized, blocked with 5% donkey serum containing 0.1% BSA and 0.1% Triton X-100, and immunostained for ZO-1 (1:200; Thermo Fisher Scientific, Cat. 339194) and VE-cadherin (1:200; Thermo Fisher Scientific, Cat. 53-1449-42) along with DAPI (1:1000; BioLegend). For live transcytosis evaluations, AF555-labeled TβR2-HFt (40 nM) was introduced into the blood-side channel of the freshly prepared chips, and transport into the brain-side channel was monitored over time by confocal microscopy (LSM 710, Carl Zeiss).

### *In vitro* 3D multicellular spheroid penetration study

3D multicellular tumor spheroids were established by the hanging-drop method as previously described (*59*). GL261-Luc2 cell suspensions (5 × 10^3^ cells in 20 μL) were cultured as hanging drops for 2 days. Spheroids were transferred to U-bottom 96-well plates (CELLSTAR, Greiner Bio-One) in RPMI 1640 medium (Thermo Fisher Scientific). Spheroids were cultured at 37 °C for 48 h until reaching approximately 500 μm in diameter. Spheroids were incubated with 10 μg of AF555-labeled HFt or TβR2-HFt for 48 h. Spheroids were transferred onto poly-D-lysine-coated glass-bottom dishes (MatTek). Z-stack images were acquired at depths up to 200 μm using confocal microscopy (LSM 710, Carl Zeiss, 10× objective).

### Flow cytometry for the assessment of TβR2-HFt binding to glioma cell surfaces

TβR2-HFt binding to glioma cell surfaces was evaluated by flow cytometry using human HSR-GBM1 cells and mouse GL261-Luc2 and CT-2A cells. Adherent cells were detached with Accutase, and HSR-GBM1 spheroids were dissociated by Accutase treatment with trituration. Single-cell suspensions were obtained using a 70-μm cell strainer. Cells (2.5 × 10^5^ cells per sample) were incubated with 1 nM AF647-labeled TβR2-HFt in ice-cold PBS containing 1% BSA for 1 h at 4 °C in the dark. Incubation at 4 °C minimized nanocage internalization. Control cells were processed under identical conditions without AF647-labeled TβR2-HFt.

Cells were washed (500 × g, 5 min, 4 °C), stained with Zombie Aqua viability dye (BioLegend), resuspended in PBS containing 1% BSA, and acquired on a Cytek Aurora flow cytometer (Cytek Biosciences). Gating was established on live singlets (FSC-A vs. FSC-H; Zombie Aqua negative) using unstained and single-stained controls. AF647 fluorescence intensity and positive cell fractions were quantified across three independent biological replicates in FlowJo (Tree Star).

### Preparation of primary orthotopic glioma and surgical resection models

All animal procedures were performed in accordance with protocols approved by the Institutional Animal Care and Use Committees of Johns Hopkins University and the University of Maryland. Orthotopic glioma models were established in 6-8-week-old female C57BL/6J mice (Jackson Laboratory) using a stereotaxic frame under anesthesia with isoflurane or ketamine/xylazine. Perioperative analgesia was administered using carprofen and local bupivacaine/lidocaine at the incision site. A burr hole was drilled 0.5 mm posterior to bregma and 2.0 mm lateral to the midline. Tumor cells (GL261-Luc2 or CT-2A; 1 × 10^5^ cells in 2 µL) were injected into the right striatum at a depth of 2.5-3.0 mm. Tumor establishment in GL261-Luc2-bearing mice was confirmed on day 7 post-implantation using an IVIS Spectrum system (PerkinElmer). Mice with bioluminescent signals exceeding tenfold background were randomized into experimental groups based on baseline flux values. Tumor establishment in CT-2A-bearing mice was confirmed on day 7 post-implantation using a 9.4T Bruker MRI system. Mice with MRI-detectable tumors were randomized into experimental groups based on baseline tumor volume.

For the preparation of a surgical resection model, tumor tissue was resected using a stereotaxic NICO Myriad NOVUS surgical aspirator equipped with a 23-gauge probe. After resection, hemostasis was achieved as needed, and the incision was closed. Residual tumor signal was assessed by bioluminescence imaging 4 h after resection. Mice with detectable post-resection bioluminescent signal above background were excluded as incomplete resections. The remaining mice were randomly assigned to treatment groups, followed by imaging and histological quantifications in a blinded manner.

### TβR2-HFt administration and *in vivo* delivery and mechanism assessments

For systemic tolerability, healthy mice were intravenously administered TβR2-HFt (2.5, 5, or 10 mg/kg) every 5 days, and body weight was monitored for 28 days (*n* = 5 per group).

The BBB penetration and access to tumor-bearing brain were assessed after intravenous injection of AF555-labeled TβR2-HFt (10 mg/kg). To evaluate brain accumulation, healthy mice (PBS, *n* = 4; AF555-labeled TβR2-HFt, *n* = 3) or GL261-Luc2 tumor-bearing mice (*n* = 3 per group) were intravenously administered PBS or AF555-labeled TβR2-HFt (10 mg/kg), and brains were harvested after PBS perfusion at 3 h and 24 h post-injection, respectively. *Ex vivo* fluorescence was imaged with an IVIS Spectrum system and quantified within defined brain regions of interest.

For mechanistic evaluations, GL261-Luc2 tumor-bearing mice received intravenous PBS or TβR2-HFt (10 mg/kg) every 5 days up to four doses. Tumor burden was assessed by bioluminescence imaging after the final dose (*n* = 5 per group). Distinct cohorts were analyzed for survival (monitored until humane endpoints) or harvested for subsequent assessments: histology (pSMAD2 IHC, *n* = 3 per group; Masson’s trichrome, *n* = 4-5 per group; Sirius Red, *n* = 3-4 per group), flow cytometry (*n* = 6-9 per group), and qPCR (*n* = 9-12 per group).

### Preparation of brain tissue sections and histological staining

Harvested mouse brains were fixed by transcardial perfusion and subsequent immersion in 10% neutral buffered formalin for 24 h, followed by standard dehydration and paraffin embedding, or cryoprotected in 30% sucrose and frozen on dry ice. Paraffin-embedded tissues were sectioned at 4 μm for pSMAD2 immunohistochemistry and at 10 μm for Masson’s trichrome and Sirius Red staining; histological staining was performed by Reference Histology at Johns Hopkins University using standard protocols and imaged on a Keyence BZ-X microscope. Cryopreserved tissues were sectioned at a 15-μm thickness on a cryostat and stored in cryoprotectant solution prior to immunofluorescence.

For pSMAD2 detection, deparaffinized and rehydrated sections were subjected to heat-induced antigen retrieval in pH 9.0 Tris-EDTA buffer, quenched for endogenous peroxidase activity, and blocked before incubation with anti-pSMAD2 antibody (1:2000; Abcam, Cat. ab280888). Signals were developed using the ImmPRESS IHC detection kit (Vector Laboratories) and counterstained with hematoxylin. For T-cell profiling, 15-μm cryosections were washed with PBS, retrieved with trypsin, permeabilized, and blocked. Sections were incubated overnight at 4 °C with anti-CD3 antibody (1:100; clone 500A2, BioLegend, Cat. 152302), anti-CD4 antibody (1:100; clone RM4-5, BioLegend, Cat. 100505), or anti-CD8 antibody (1:100; clone 53-6-7, BioLegend, Cat. 100701), followed by species-appropriate fluorescent secondary antibodies, mounting in ProLong Glass Antifade Mountant with NucBlue (Invitrogen), and curing for 48 h in the dark.

### Image-based histological analysis

Collagen-positive areas in Masson’s trichrome-stained sections were quantified in Fiji (ImageJ, NIH) using a color deconvolution method adapted from a previous study (*61*), with manual threshold parameters set to hue 135-195, saturation 25-255, and brightness 20-255. For Sirius Red and pSMAD2 staining, RGB images were converted to 8-bit grayscale, and the percentage of positive area relative to total image area was determined using threshold cutoffs of ≥195 and ≥200, respectively.

All analyses were conducted in a blinded manner using identically sized regions of interest (ROIs) within histologically defined tumor margins and uniform thresholding parameters across groups. For collagen and pSMAD2 analyses, values from multiple ROIs per tumor were averaged to yield a single value per animal.

For CD3, CD4, and CD8 immunostaining, non-consecutive tumor sections (*n* = 12 sections from 3 mice per group) were analyzed to evaluate the spatial distribution of T cells. Positive cell densities were quantified across five randomly selected tumor ROIs per section and normalized to tissue area (cells per mm²), with intra-section ROI measurements averaged to generate individual data points for statistical evaluation.

### Brain tissue preparation for the analysis of mRNA transcript levels

For baseline analysis, brains from untreated mice with IVIS-confirmed GL261-Luc2 tumors (*n* = 3) were excised following PBS perfusion, and the tumor-bearing and contralateral hemispheres were collected. To evaluate therapy-induced microenvironmental changes, mice receiving PBS or TβR2-HFt were euthanized the day after the fourth administration and perfused with PBS. The anterior half of the tumor-injected right cerebral hemisphere was collected as tumor and peritumoral tissue, alongside corresponding brain regions from tumor-free healthy mice. Harvested brain tissues were immediately snap-frozen in liquid nitrogen, pulverized, and homogenized using a TissueLyser LT (Qiagen). Total RNA isolation and RT-qPCR were performed as described below.

### Immune profiling of brain tumor tissues by flow cytometry

One day after the fourth dose, PBS-perfused brain tumors were harvested, minced, and digested with Collagenase D (Roche, Cat. 11088882001) for 1 h at 37 °C as previously described (*62*). Digested tissues were passed through a 70-μm strainer (Fisher Scientific, Cat. 22-363-548), treated with RBC Lysis Buffer (BioLegend, Cat. 420301), and subjected to discontinuous 40%/80% Percoll density-gradient centrifugation (Cytiva, Cat. 17089101) to enrich tumor-infiltrating immune cells. Enriched leukocytes were stained with Zombie Aqua viability dye and surface antibodies for 30 min at 4 °C. All recoverable events (≥5,000 events per sample; typically >10,000 events) were acquired on a Cytek Aurora 3 spectral cytometer (Cytek Biosciences) and analyzed in FlowJo.

For intracellular cytokine profiling, CD8^+^ T cells were isolated from tumor single-cell suspensions using the MojoSort™ Mouse CD8 T Cell Isolation Kit (BioLegend, Cat. 480008) and split into paired unstimulated or restimulated conditions. Restimulation was conducted using Cell Activation Cocktail with Brefeldin A (2 μL/mL; BioLegend, Cat. 423303) for 6 h at 37 °C in 5% CO₂, followed by surface CD8 and intracellular IFN-γ staining.

Gating was initiated on FSC-A versus SSC-A, followed by singlet discrimination (FSC-A vs. FSC-H) and dead-cell exclusion (Zombie Aqua). Within live CD3^+^CD11b^−^ T cells, subsets were delineated as CD4^+^ Treg cells (CD4^+^ CD25^+^) and effector CD8^+^ T cells (CD8^+^ PD-1^int^ TIM-3^−^), and quantified as frequencies within total CD3*^+^*T cells. For IFN-γ analysis, live singlet CD3^+^ CD8^+^ T cells were gated from the isolated CD8^+^ fraction, and IFN-γ^+^ frequencies were established using matched unstimulated controls. Positive gates were set using single-stain and unstained autofluorescence controls and applied uniformly across experiments. Antibodies and representative gating hierarchies are provided in **Table S5** and **Fig. S9**, respectively.

### *In vitro* human glioma spheroid and astrocyte studies

HSR-GBM1 spheroids were cultured in 6-well plates. Human astrocytes were differentiated from H9 hESCs via neural induction using the STEMdiff SMADi Neural Induction Kit (STEMCELL Technologies), followed by sequential 3-week differentiation and 3-week maturation using STEMdiff Astrocyte Differentiation and Serum-Free Maturation Kits, respectively. Cells were assigned to three treatment cohorts: vehicle control, hTGFβ1, and TβR2-HFt plus hTGFβ1. For the combination cohort, cells were pretreated with 1 µg/mL TβR2-HFt for 5 min prior to 5 ng/mL hTGFβ1 stimulation. Control cohorts received 5 ng/mL hTGFβ1 alone or matched vehicle volumes. After 24 h of incubation at 37 °C and 5% CO₂, cells were harvested for RNA isolation. Independent biological replicates were analyzed for HSR-GBM1 spheroids (*n* = 6 per group) and astrocytes (*n* = 3-4 per group). Total RNA isolation and RT-qPCR analysis were performed as described below.

### RNA isolation and RT-qPCR

Total RNA was isolated using the RNeasy Plus Mini Kit (Qiagen), reverse-transcribed (High-Capacity cDNA RT Kit, Thermo Fisher Scientific), and analyzed via qPCR with PowerTrack SYBR Green Master Mix on a QuantStudio 3 system (Applied Biosystems). Relative mRNA transcript levels were determined in technical triplicate using the 2^−ΔΔCt^ method normalized to *Gapdh* (mouse) or *GAPDH* (human) using primers in **Table S6**. Samples lacking reliable Ct values (Ct > 35 or undetermined amplification) were excluded.

### Therapeutic efficacy assessments

An independent cohort of GL261-Luc2 tumor-bearing mice with IVIS-confirmed tumors on day 7 post-implantation were randomized into four groups (*n* = 9 per group): PBS, anti-PD-1 mAb monotherapy (10 mg/kg; clone RMP1-14; BioXCell, Cat. BE0146), anti-PD-1 mAb (10 mg/kg) plus anti-TGFβ mAb (10 mg/kg; clone 1D11.16.8; BioXCell, Cat. BE0057), or anti-PD-1 mAb plus TβR2-HFt (10 mg/kg). Doses were selected based on prior studies (*10, 60*). All agents were administered intravenously at 10 mg/kg every 5 days from day 7 post-implantation until humane endpoints were reached.

For the CT-2A orthotopic model study, mice with MRI-confirmed tumors on day 7 were randomized based on baseline tumor volume into three groups (*n* = 9-10 per group): PBS, anti-PD-1 mAb monotherapy, or anti-PD-1 mAb plus TβR2-HFt. Starting on the day of MRI confirmation, treatments were intravenously given every 2 days up to three doses, and mice were monitored until humane endpoints.

For the resection model study, beginning on the day of surgery, mice received intravenous PBS, anti-PD-1 mAb, or anti-PD-1 mAb plus TβR2-HFt every 5 days up to three doses. Postoperative tumor growth was monitored by bioluminescence imaging. Mice were monitored until humane endpoints were reached and were recorded as deceased at the time of euthanasia.

Long-term survival mice from the resection study were rechallenged on day 120 post-resection via contralateral orthotopic implantation of GL261-Luc2 cells (1 × 10^5^ cells in 2 µL). Age-matched healthy control mice were identically inoculated for comparison. Mice were subsequently monitored without additional treatment for 90 days or until reaching humane endpoints.

### Statistical analysis

Data are presented as mean ± s.d. unless otherwise indicated, and analyses were performed in GraphPad Prism (v10.2.3). For *in vitro* assays, experiments were performed in independent biological triplicates unless otherwise indicated, and technical replicates were averaged prior to statistical analysis. Two-group comparisons were evaluated using two-tailed unpaired Student’s *t*-tests, or paired *t*-tests for matched tumor-bearing and contralateral hemispheres. Multi-group comparisons were analyzed by one-way ANOVA with Tukey’s post hoc test or two-way repeated-measures ANOVA with Sidak’s multiple comparisons test where indicated. Survival curves were assessed using the log-rank test with Bonferroni correction for multiple cohorts, and correlations were evaluated by Pearson’s correlation analysis. ns, not significant; \**p* < 0.05, \*\**p* < 0.01, \*\*\**p* < 0.001, \*\*\*\**p* < 0.0001.

## Data availability

The data that support the findings of this study are available from the corresponding author upon reasonable request.

## Acknowledgements

The funding was provided by the National Institutes of Health (R01NS119609, R01NS133965, and P30EY001765). This research was also supported by the Korea Health Industry Development Institute under the Ministry of Health and Welfare of the Republic of Korea (Korea Health Technology R&D Project; HI19C0642).

We thank Haolin Li and Kijung Kwak for their assistance with preliminary large-scale animal studies. We also thank Zixu Han and Sri Sai Vivek Alla for their assistance with recombinant protein production. We are especially grateful to Dr. Gijung Kwak for his assistance with preliminary large-scale animal experiments as well as insightful discussion.

## Author contributions

J. S. Suk conceived and supervised the study, acquired funding, and revised the manuscript. D. Lee designed and performed experiments, analyzed and interpreted data, curated figures, and wrote the manuscript. S. W. Chung performed experiments and contributed to experimental methodologies and data acquisition. B. Kong contributed to experiments, data acquisition, and figure preparation. Y. Xie, B. Kang, J. Yang, B. Huang, D. Rao, J. S. Choi, and P. Anastasiadis contributed to experiments and data acquisition. E. A. Ra prepared human cell resources. G. Lee and D.-H. Kim provided scientific support and relevant expertise. M. Lim provided scientific support and expertise in immunotherapy as well as contributed to study discussions. All authors approved the final manuscript.

