## Supplementary Information for "Blood-brain barrier-traversing multivalent TGFβ trap for anti-cancer immunomodulation of gliomas"

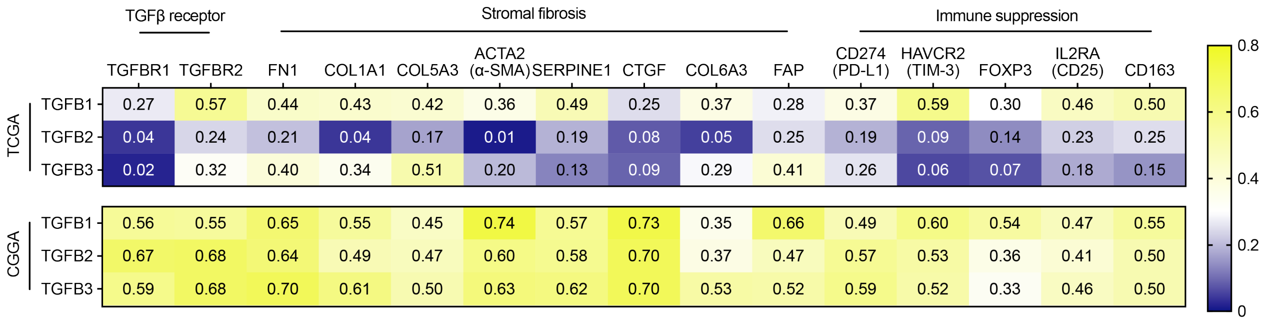


**Fig. S1. TGFβ isoform levels positively correlate with stromal fibrosis and immune suppression markers in TME of TCGA-GBM and CGGA-GBM cohorts.** Pearson's correlation coefficients showing the correlations between *TGFB1*, *TGFB2*, and *TGFB3* mRNA transcript levels and those of TGFβ receptors, stromal fibrosis markers, or immune suppression markers in TCGA-GBM (top) and CGGA-GBM (bottom) cohorts. TGFβ receptors: *TGFBR1* and *TGFBR2*; Stromal fibrosis markers: *FN1*, *COL1A1*, *COL5A3*, *ACTA2*, *SERPINE1*, *CTGF*, *COL6A3*, and *FAP*; Immune suppression markers: *CD274*, *HAVCR2*, *FOXP3*, *IL2RA*, and *CD163*.

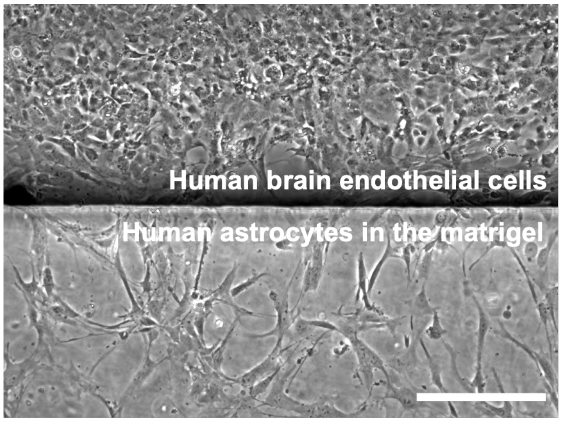


**Fig. S2. A physically distinct barrier is established in a 3D microfluidic BBB chip model based on human brain endothelial cells and human astrocytes.** Phase-contrast image showing a well-defined boundary between the blood-side channel and the brain-side channel. Scale bar, 50 μm.

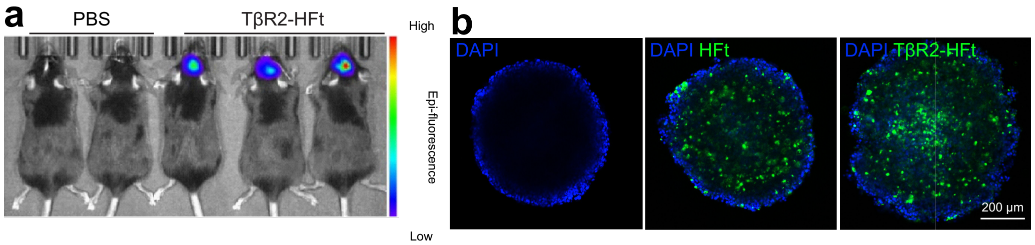


**Fig. S3. TβR2-HFt accumulates in orthotopic mouse glioma-bearing brains *in vivo* following systemic administration and penetrates 3D mouse glioma spheroids *in vitro*. a**, Live-animal IVIS image showing accumulation of fluorescently labeled TβR2-HFt in mouse brains bearing orthotopic GL261-Luc2 tumors. **b**, Representative confocal images showing distribution of fluorescently labeled HFt and TβR2-HFt within 3D multicellular tumor spheroids based on GL261-Luc2 cells. Blue: DAPI; green: HFt/TβR2-HFt, Scale bars, 200 μm.

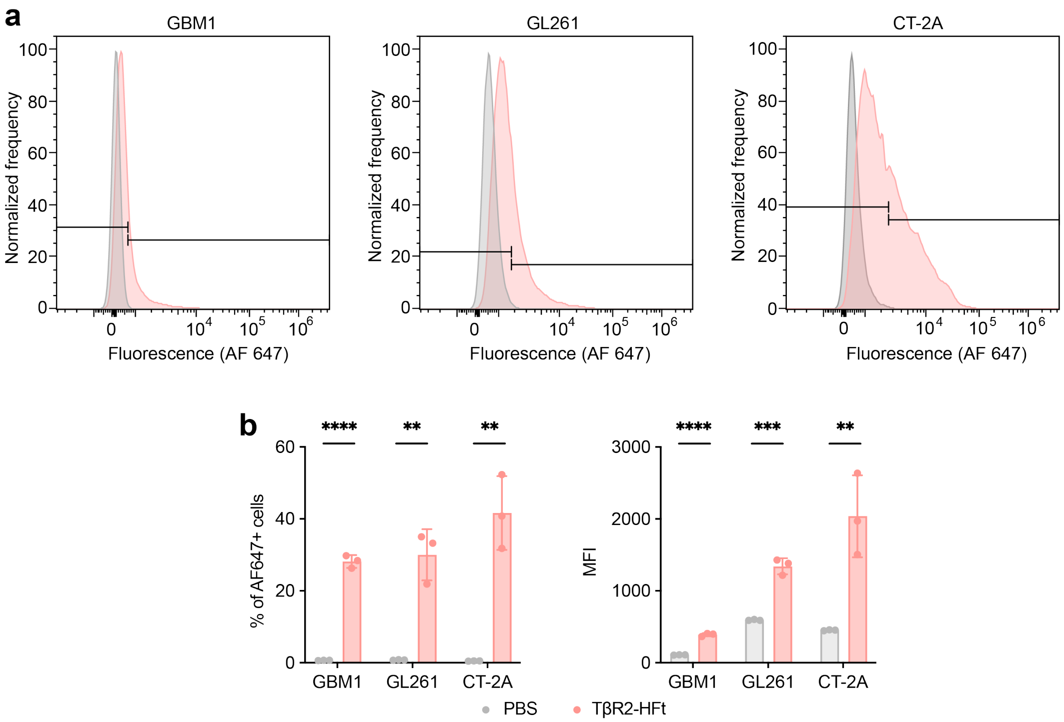


**Fig. S4. TβR2-HFt binds to human and mouse glioma cell surfaces. a,** Representative flow cytometry histograms showing AF647 fluorescence signals associated with human HSR-GBM1 (left) and mouse GL261-Luc2 (middle) and CT-2A (right) glioma cells after incubation with AF647-labeled TβR2-HFt at 4 °C to minimize internalization. Untreated cells were used as a negative control. **b,** Frequency of AF647-positive cells and median fluorescence intensity measured for untreated and AF647-TβR2-HFt-treated cells shown in a (n = 3). Data are presented as mean ± s.d. Statistical significance was determined by unpaired two-tailed Student's t-test. $^{**}p < 0.01$, $^{***}p < 0.001$, $^{****}p < 0.0001$.

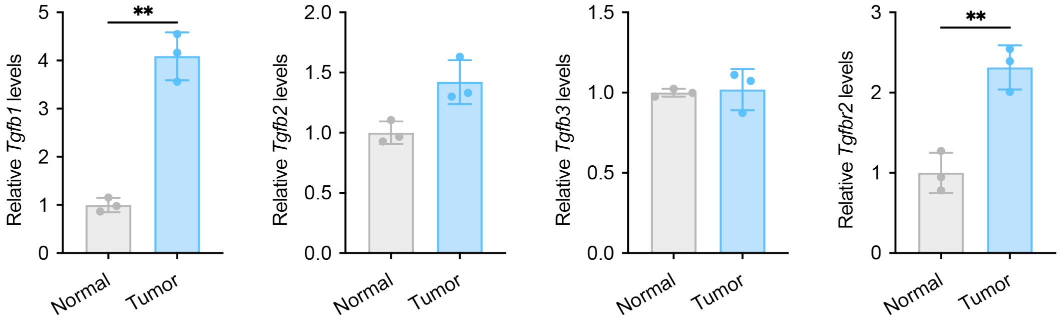


**Fig. S5. Orthotopic mouse gliomas show *Tgfb1* and *Tgfbr2* upregulation.** Relative *Tgfb1*, *Tgfb2*, *Tgfb3*, and *Tgfbr2* mRNA transcript levels in tumor-bearing and contralateral tumor-free hemispheres of mouse brains, measured by qPCR. Data are presented as mean ± s.d. Statistical significance was determined by paired two-tailed Student's t-test. *$p < 0.05$, **$p < 0.01$. $p = 0.0824$ for *Tgfb2*.

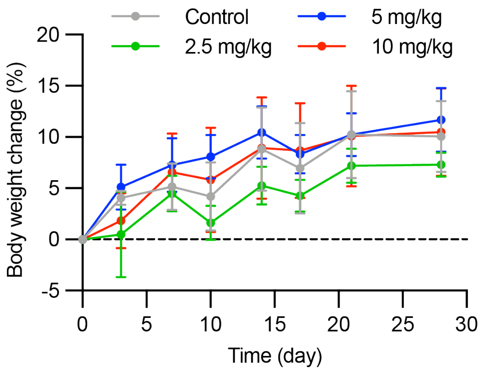


**Fig. S6. Repeated systemic administration of TβR2-HFt does not cause body-weight loss of healthy mice.** Body weight changes in healthy C57BL/6J mice following systemic administration of PBS or TβR2-HFt at a dose of 2.5, 5, or 10 mg/kg every 5 days for 28 days. Data are presented as mean ± s.d.

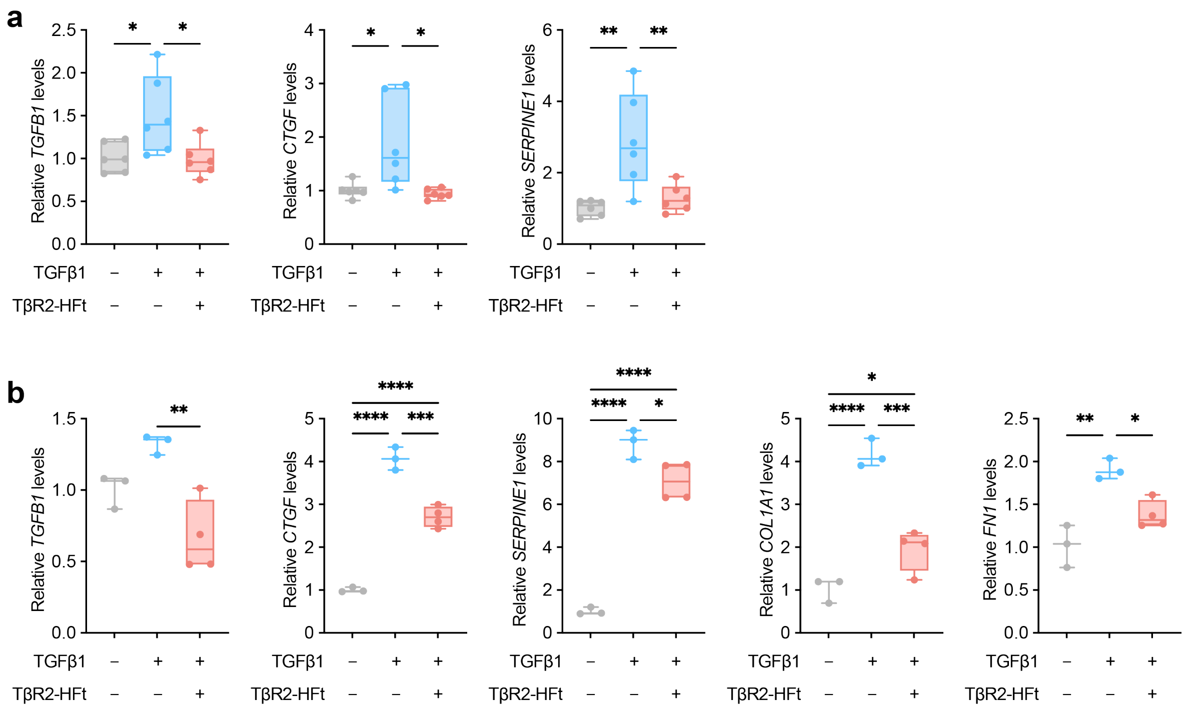


**Fig. S7. TβR2-HFt suppresses TGFβ-induced profibrotic programs in human HSR-GBM1 cells and human astrocytes. a**, Relative levels of *TGFB1*, *CTGF*, and *SERPINE1* mRNA transcripts in HSR-GBM1 cells treated with recombinant human TGFβ1 in presence or absence of TβR2-HFt. **b**, Relative levels of *TGFB1*, *CTGF*, *SERPINE1*, *COL1A1*, and *FN1* mRNA transcripts in H9 hESC-derived human astrocytes treated with recombinant human TGFβ1 in presence or absence of TβR2-HFt. Boxes, center lines, and whiskers indicate interquartile range, median, and minimum and maximum values, respectively (HSR-GBM1 spheroid, n = 6; astrocyte, n = 3 - 4). Statistical significance was determined by one-way ANOVA with Tukey's post hoc test. *$p < 0.05$, **$p < 0.01$, ***$p < 0.001$, ****$p < 0.0001$.

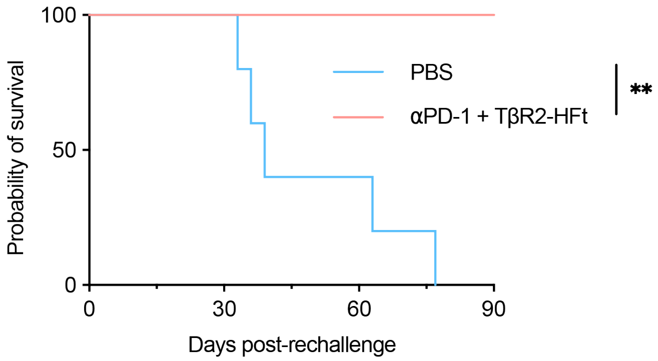


**Fig. S8. All long-term survival animals resist orthotopic GL261-Luc2 tumor rechallenge without additional treatment.** Kaplan-Meier survival analysis after contralateral orthotopic GL261-Luc2 rechallenge of long-term survival animals from the post-resection anti-PD-1 plus TβR2-HFt treatment group in Fig. 7e in comparison to the identically tumor-inoculated age-matched naive control mice (n = 5 per group). Long-term survival animals were rechallenged on day 120 after surgical resection and monitored for additional 90 days without additional treatment. Statistical significance was determined by the log-rank test. $p = 0.0054$.

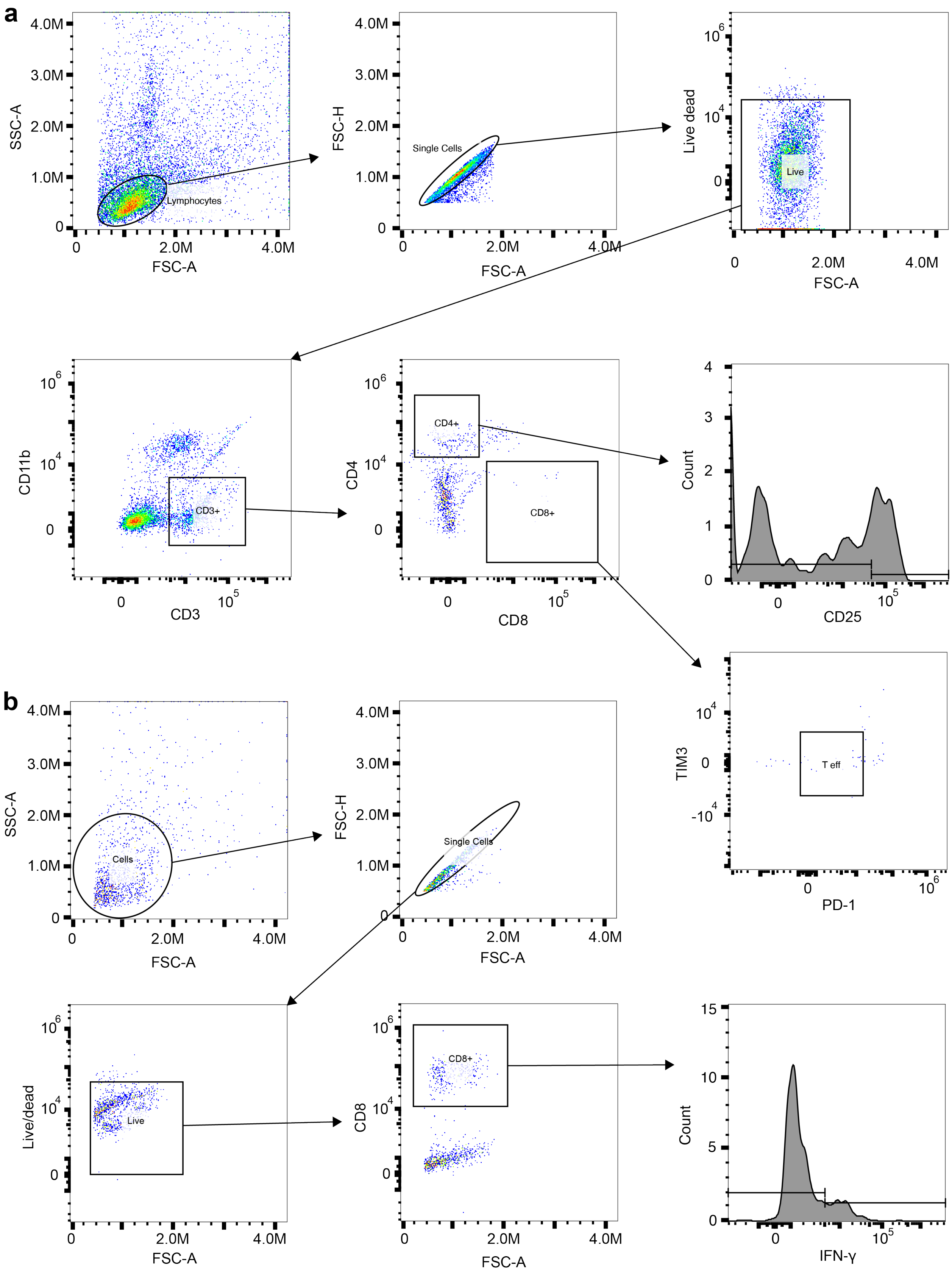
a
SSC-A
FSC-A
Lymphocytes
FSC-H
Single Cells
Live dead
Live
CD11b
CD3
CD3+
CD4
CD8
CD4+
CD8+
Count
CD25
TIM3
PD-1
T eff
b
Cells
Live/dead
Live
IFN-γ

**Fig. S9. Representative flow cytometry gating strategy. a,** Gating strategy for tumor-infiltrating lymphocyte analysis. Total lymphocytes were initially identified by FSC-A versus SSC-A, followed by doublet discrimination using FSC-A versus FSC-H (Single Cells) and dead-cell exclusion using Zombie Aqua viability dye (Live). $CD3^+$ T cells were then gated and subsequently separated into $CD4^+$ and $CD8^+$ T-cell subsets. Regulatory T cells were defined as $CD25^+$ cells within the $CD4^+$ population (histogram). Effector (T eff, PD-$1^{int}$ $TIM3^-$) and exhausted (T ex, PD-$1^{high}$ $TIM3^+$) $CD8^+$ T-cell populations were identified from the CD8+ gate using PD-1 versus TIM-3. **b,** Gating strategy for intracellular IFN-γ analysis. Cells isolated from brain tumors were gated by FSC-A versus SSC-A, followed by singlet gating (FSC-A versus FSC-H) and dead-cell exclusion. $CD8^+$ T cells were gated, and the IFN-$\gamma^+$ fraction was quantified via histogram analysis following *ex vivo* re-stimulation. Gates were established using matched unstimulated and single-stained controls.

**Table S1. Amino acid sequences of fusion protein construct variants.** Blue: FHC; red: TGFβR2ecd.

| Variant | Amino acid sequence |
|---|---|
| 1 | HHHHHHENLYFQGTTASTSQVRQNYHQDSEAAINRQINLELYASYVYLSMSYY<br>FDRDDVALKNFAKYFLHQSHEEREHAEKLMKLQNQRGGRIFLQDIKKPDCDD<br>WESGLNAMECALHLEKNVNQSLLELHKLATDKNDPHLCDFIETHYLNEQVKAI<br>KELGDHVTNLRKMGAPESGLAEYLFDKHTLGDSDNESGGGGSGGGGSGGGGS<br>GGGGSTIPPHVQKSVNNDMIVTDNNGAVKFPQLCKFCDVRFSTCDNQKSCMSN<br>CSITSICEKPQEVCVAVWRKNDENITLETVCHDPKLPYHDFILEDAASPKCIMKE<br>KKKPGETFFMCSCSSDECNDNIIFSEEYNTSNPD |
| 2 | HHHHHHENLYFQGTTASTSQVRQNYHQDSEAAINRQINLELYASYVYLSMSYY<br>FDRDDVALKNFAKYFLHQSHEEREHAEKLMKLQNQRGGRIFLQDIKKPDCDD<br>WESGLNAMECALHLEKNVNQSLLELHKLATDKNDPHLCDFIETHYLNEQVKAI<br>KELGDHVTNLRKMGAPESGLAEYLFDKHTLGDSDNESAEAAAKEAAAKEAAA<br>KEAAAKALEAEAAAKEAAAKEAAAKEAAAKATIPPHVQKSVNNDMIVTDNNG<br>AVKFPQLCKFCDVRFSTCDNQKSCMSNCSITSICEKPQEVCVAVWRKNDENITL<br>ETVCHDPKLPYHDFILEDAASPKCIMKEKKKPGETFFMCSCSSDECNDNIIFSEE<br>YNTSNPD |
| 3 | HHHHHHENLYFQGTIPPHVQKSVNNDMIVTDNNGAVKFPQLCKFCDVRFSTCD<br>NQKSCMSNCSITSICEKPQEVCVAVWRKNDENITLETVCHDPKLPYHDFILEDA<br>ASPKCIMKEKKKPGETFFMCSCSSDECNDNIIFSEEYNTSNPDGGGGSGGGGSG<br>GGGSGGGGSTTASTSQVRQNYHQDSEAAINRQINLELYASYVYLSMSYYFDRD<br>DVALKNFAKYFLHQSHEEREHAEKLMKLQNQRGGRIFLQDIKKPDCDDWESGL<br>NAMECALHLEKNVNQSLLELHKLATDKNDPHLCDFIETHYLNEQVKAIKELGD<br>HVTNLRKMGAPESGLAEYLFDKHTLGDSDNES |
| 4 | HHHHHHENLYFQGTIPPHVQKSVNNDMIVTDNNGAVKFPQLCKFCDVRFSTCD<br>NQKSCMSNCSITSICEKPQEVCVAVWRKNDENITLETVCHDPKLPYHDFILEDA<br>ASPKCIMKEKKKPGETFFMCSCSSDECNDNIIFSEEYNTSNPDAEAAAKEAAAK<br>EAAAKEAAAKALEAEAAAKEAAAKEAAAKEAAAKATTASTSQVRQNYHQDS<br>EAAINRQINLELYASYVYLSMSYYFDRDDVALKNFAKYFLHQSHEEREHAEKL<br>MKLQNQRGGRIFLQDIKKPDCDDWESGLNAMECALHLEKNVNQSLLELHKLA<br>TDKNDPHLCDFIETHYLNEQVKAIKELGDHVTNLRKMGAPESGLAEYLFDKHT<br>LGDSDNES |

**Table S2. The Kd values (pM) calculated based on the binding curves in Fig. 4a.**

| | TGFβ1 | TGFβ2 | TGFβ3 |
|---|---|---|---|
| TβR2-HFt | 5.579 (4.921–6.330) | 26.68 (23.68–30.13) | 45.42 (39.33–52.51) |
| Pan-TGFβ mAb (1D11) | 10.79 (10.41–11.19) | 30.00 (24.99–36.27) | 96.84 (61.39–181.9) |

The ranges in parentheses are 95% confidence intervals.

**Table S3. The Kd values (pM) calculated based on the binding curves in Fig. 5b.**

| | hTfR1 | mTIM-2 |
|---|---|---|
| HFt | 352.3 (317.5–390.6) | 6.839 (5.744–8.128) |
| TβR2-HFt | 817.0 (679.2–980.7) | 19.95 (17.28–23.01) |

The ranges in parentheses are 95% confidence intervals.

**Table S4. Pairwise survival comparisons in orthotopic and post-resection glioma models.**

| Model | Group 1 | Group 2 | Median survival, Group 1 | Median survival, Group 2 | Raw p-value | Adjusted p-value |
|---|---|---|---|---|---|---|
| GL261 | PBS | Anti-PD-1 | 30 | 38 | 0.0551 | 0.3306 |
| | PBS | Anti-PD-1 + 1D11 | 30 | 41 | 0.0763 | 0.4578 |
| | PBS | Anti-PD-1 + TβR2-HFt | 30 | 47 | 0.0006 | 0.0036 |
| | Anti-PD-1 | Anti-PD-1 + 1D11 | 38 | 41 | 0.7188 | > 0.9999 |
| | Anti-PD-1 | Anti-PD-1 + TβR2-HFt | 38 | 47 | 0.0227 | 0.1362 |
| | Anti-PD-1 + 1D11 | Anti-PD-1 + TβR2-HFt | 41 | 47 | 0.0248 | 0.1488 |
| CT-2A | PBS | Anti-PD-1 | 17 | 19 | 0.3715 | > 0.9999 |
| | PBS | Anti-PD-1 + TβR2-HFt | 17 | 27 | 0.0047 | 0.0141 |
| | Anti-PD-1 | Anti-PD-1 + TβR2-HFt | 19 | 27 | 0.0350 | 0.1050 |
| Resection | PBS | Anti-PD-1 | 40 | 46 | 0.1699 | 0.5097 |
| | PBS | Anti-PD-1 + TβR2-HFt | 40 | not reached | 0.0160 | 0.0480 |
| | Anti-PD-1 | Anti-PD-1 + TβR2-HFt | 46 | not reached | 0.3180 | 0.9540 |

Pairwise survival comparisons were performed using the log-rank test. Adjusted p-values were calculated using Bonferroni correction within each experiment.

**Table S5**. **Antibodies and viability dye used for flow cytometry.**

| Target | Fluorophore | Clone | Vendor | Catalog number |
|---|---|---|---|---|
| Fixable viability dye | Zombie Aqua | N/A | BioLegend | 423101 |
| CD11b | PE-Dazzle | M1/70 | BioLegend | 128049 |
| CD3 | Brilliant Violet 711 | 17A2 | BioLegend | 100241 |
| CD4 | Brilliant Violet 570 | RM4-5 | BioLegend | 100542 |
| CD8α | PE/Cyanine5 | 53-6.7 | BioLegend | 100710 |
| CD25 | PE | PC61 | BioLegend | 102008 |
| TIM3 / CD366 | Brilliant Violet 785 | RMT3-23 | BioLegend | 119725 |
| PD-1 / CD279 | APC | 29F.1A12 | BioLegend | 135210 |
| IFN-γ | BV421 | XMG1.2 | BioLegend | 505830 |

**Table S6**. **Primer sequences used for qPCR.**

| Species | Gene symbol | Forward | Reverse |
|---|---|---|---|
| Mouse | *Tgfb1* | TGATACGCCTGAGTGGCTGTCT | CACAAGAGCAGTGAGCGCTGAA |
| | *Tgfb2* | TTGTTGCCCTCCTACAGACTGG | GTAAAGAGGGCGAAGGCAGCAA |
| | *Tgfb3* | AAGCAGCGCTACATAGGTGGCA | GGCTGAAAGGTGTGACATGGAC |
| | *Tgfbr2* | CCTACTCTGTCTGTGGATGACC | GACATCCGTCTGCTTGAACGAC |
| | *Col1a1* | CCTCAGGGTATTGCTGGACAAC | CAGAAGGACCTTGTTTGCCAGG |
| | *Col5a3* | GGCAAAGATGGTATTCCAGGACC | TGCTTCCTTTGTGACCAGGCATC |
| | *Fn1* | CCCTGTTCTGCTTCAGGGTT | AAAGCAGAGGTGTCTGGGTG |
| | *Acta2* | GTACGTGTTGATCCTGACTGCTGTCATGG | TCATTTCTTCCTGGGCTGCTTATCTGG |
| | *Gapdh* | CATCACTGCCACCCAGAAGACTG | ATGCCAGTGAGCTTCCCGTTCAG |
| Human | *TGFB1* | CAATTCCTGGCGATACCTCAG | GCACAACTCCGGTGACATCAA |
| | *SERPINE1* | ACCGCAACGTGGTTTTCTCA | TTGAATCCCATAGCTGCTTGAAT |
| | *CTGF* | CAGCATGGACGTTCGTCTG | AACCACGGTTTGGTCCTTGG |
| | *FN1* | AGGAAGCCGAGGTTTTAACTG | AGGACGCTCATAAGTGTCACC |
| | *COL1A1* | GTGCGATGACGTGATCTGTGA | CGGTGGTTTCTTGGTCGGT |
| | *GAPDH* | ACAACTTTGGTATCGTGGAAGG | GCCATCACGCCACAGTTTC |
